# Alveolar epithelial cell plasticity and injury memory in human pulmonary fibrosis

**DOI:** 10.1101/2025.06.10.658504

**Authors:** Taylor S Adams, Jonas C Schupp, Agshin Balayev, Johad Khoury, Aurélien Justet, Fadi Nikola, Laurens J De Sadeleer, Juan Cala-Garcia, Marta Zapata-Ortega, Panayiotis V Benos, John E McDonough, Farida Ahangari, Melanie Königshoff, Jun Ding, Robert J Homer, Ivan Rosas, Xiting Yan, Bart M Vanaudenaerde, Wim A Wuyts, Naftali Kaminski

**Author notes:** **Corresponding author:** Naftali Kaminski. Boehringer-Ingelheim Endowed Professor of Internal Medicine. Chief of Pulmonary, Critical Care and Sleep Medicine. Yale School of Medicine. 300 Cedar Street, TAC– 441 South. P.O. Box 208057. New Haven, CT 06520-8057. Both authors contributed equally to this work (Taylor S Adams, Jonas C Schupp).

## Abstract

Acute and repetitive lung epithelial injury can lead to irreversible and even progressive pulmonary fibrosis; Idiopathic pulmonary fibrosis (IPF) is a fatal disease and quintessential example of this phenomenon. The composition of epithelial cells in human pulmonary fibrosis – irrespective of disease etiology – is marked by the presence of Aberrant Basaloid cells: an abnormal cell phenotype with pro-fibrotic and senescent features, localized to the surface of fibrotic lesions. Despite their relevance to human pulmonary fibrosis, the exotic molecular profile of Aberrant Basaloid cells has obscured their etiology, preventing insights into how or why these cells emerge with fibrosis. Here we identify cellular intermediaries between Aberrant Basaloid and normal alveolar epithelial cells in human IPF tissue. We track the emergence of Aberrant Basaloid cells from alveolar epithelial cells *ex vivo* and uncover a role for similar cells in epithelial regeneration under normal conditions. Lastly, we characterize the epigenetic changes that distinguish Aberrant Basaloid cells from their progenitors and identify hallmarks of AP-1 injury memory retention. This study elucidates the phenomenon of maladaptive epithelial plasticity and regeneration in pulmonary fibrosis and re-contextualizes therapeutic strategies for epithelial dysfunction.

## Main

The alveoli of the lung are the primary interface between the atmosphere and our circulation. Each alveolus is structurally optimized to maximize two critical functions: gas exchange and pulmonary compliance. On the alveolar surface, these tasks are delegated to two distinct types of epithelial cells: alveolar epithelial type I (ATI) cells enable gas-diffusion, while alveolar type II (ATII) cells produce surfactants that reduce surface tension^1,2^. Environmental exposure carries the risk of injury and ATII serve as the primary stem cells for alveolar epithelial regeneration. While alveolar epithelial injury and regeneration are a normal part of life, repeated injury can potentially lead to progressive and irreversible fibrosis^3^. Understanding both functional and maladaptive responses to alveolar epithelial regeneration remain critical topics in pulmonary medicine^4,5^.

Single cell RNA sequencing (scRNAseq) studies enabled the discovery of Aberrant Basaloid cells in tissue from patients with end-stage human lung fibrosis^6,7^. Aberrant Basaloid cells lack hallmarks of ATII and ATI cells and are characterized by a pronounced epithelial mesenchymal transition (EMT) signal (CDH2, VIM, COL1A1) and some overlapping features with airway basal cells (KRT17, TP63, LAMB3). Notably, this Aberrant Basaloid cell phenotype is found across many diseases including interstitial lung diseases like idiopathic pulmonary fibrosis (IPF)^6,7^ and systemic sclerosis^8^, COPD^7^ and COVID-19 induced fibrosis^9,10^. A concurrent discovery was made in mice of Transitional Krt8+ cells: a cell phenotype which emerges from ATII following pulmonary injury and persist until resolution before becoming ATI cells^11–16^. Murine Transitional Krt8+ cells are a conserved phenotype following genotoxic, endotoxic and mechanical forms of lung injury: they lack hallmarks of ATII and ATI cells and are characterized by expression of Cldn4, Krt8, Krt19, Sfn and Fn1^11–16^. Despite their situational likeness, murine Transitional Krt8+ cells lack most defining features of human disease Aberrant Basaloid cells^17^; this lack of apparent molecular similarity raises questions regarding the degree of cellular orthology between the two epithelial phenotypes. Unlike Transitional Krt8+ cells, the cellular origin of Aberrant Basaloid cells in human disease remains unknown. More importantly, it remains unclear whether Aberrant Basaloid cells play a normal role in human epithelial injury repair or whether their extreme expression profile and persistence through end-stage fibrosis are indicative of an abnormal phenomenon.

In this study, we perform single nuclei RNA sequencing (snRNAseq) on differentially fibrotic regions in human IPF lungs and identify intermediate states between Aberrant Basaloid cells and other epithelial cell types. We longitudinally profile a model of Aberrant Basaloid development and uncover a role for these cells in normal regeneration. Lastly, we dissect the epigenetic profile of Aberrant Basaloid cells and describe what makes them fundamentally different from normal epithelial cells.

## Results

Tissue specimens of differentially fibrotic IPF and control lungs were prepared and evaluated as described by McDonough *et al.*^18^. Explanted IPF lungs or donor lungs declined for transplantation were inflated, frozen and CT scanned. Small cores of tissue were isolated and subject to microCT assessment of surface density: a proxy of localized fibrosis (Fig. 1A). snRNAseq was then performed on 54 control and IPF cores with variable levels of fibrosis, and cell types were classified into major lineages (Figures 1A, S1A). To provide further context, epithelial nuclei were integrated with epithelial cell data from two previously published scRNAseq datasets of lung disease (Fig. 1A; S1A)^6,7^ before cell types were collectively re-classified (Fig. 1B, Supp. Table S1).

**Figure 1:**
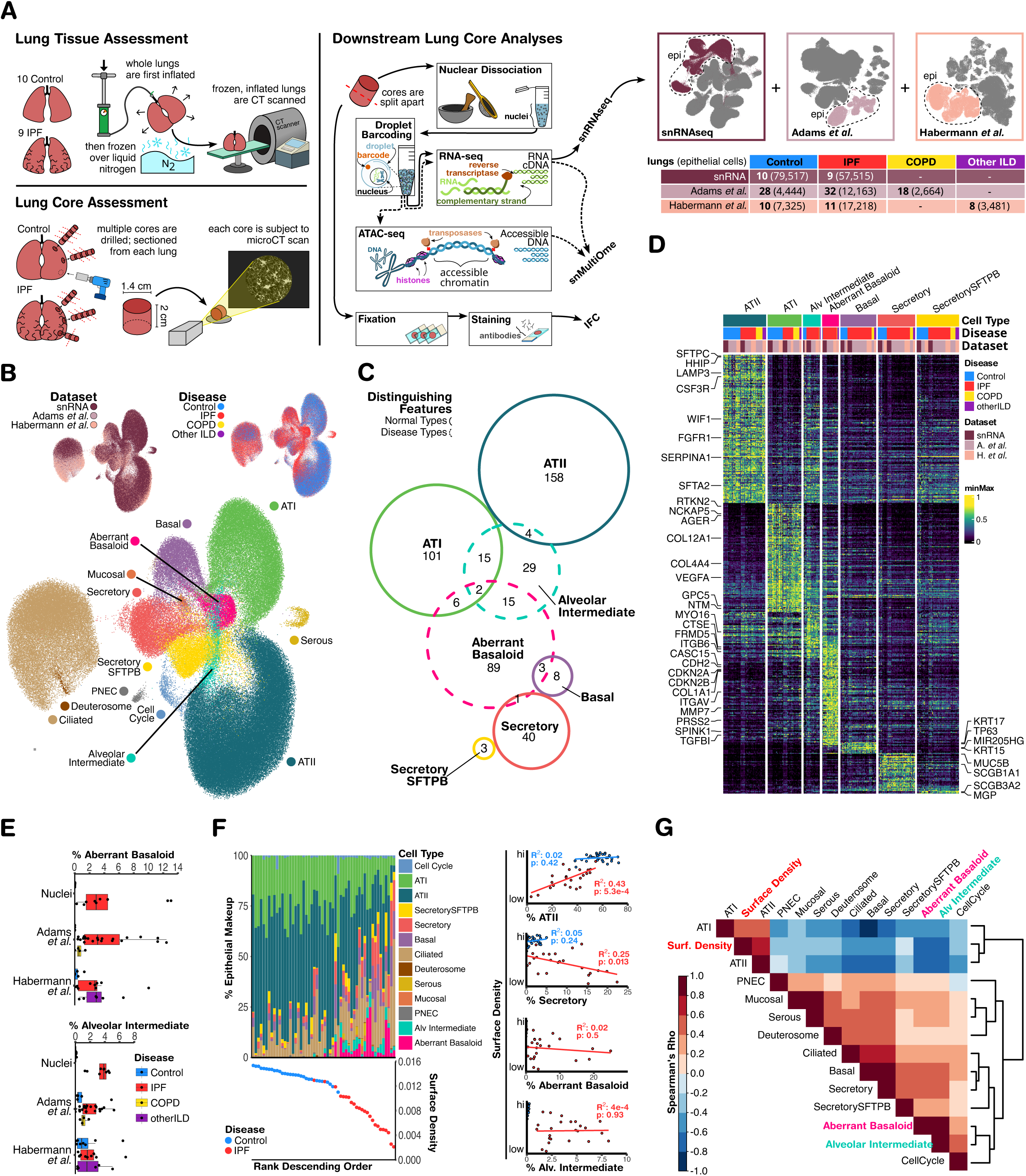
Study design and epithelial cell atlas of interstitial lung disease. A) Overview of tissue procurement and experiment design. B) UMAPs of epithelial cells from integrated analysis of multiple datasets. C) Euler diagram of intersecting conserved gene expression markers between normal and disease-associated cell types. D) Heatmap of corresponding features. Each column represents the average normalized gene expression of one patient. Within each dataset, values for each gene are min-max normalized to a common range. E) Boxplots of cell type abundances normalized to the total number of epithelial cells per subject. F) Left: Stacked bar plots of epithelial cell compositions per core in descending rank order of tissue surface density. Right: Linear regression models of changes in epithelial cell proportions as a function of surface density. G) Heatmap of spearman distances among cell-type proportions and surface density, across lung cores.

### A “Missing Link” Between Aberrant Basaloid and Alveolar Epithelial Cells

The unbiased cell representation afforded by snRNAseq (Figure S1A) enabled us to identify a previously overlooked population of epithelial cells in IPF, herein referred to as Alveolar Intermediate (AT*i*) cells (Figure 1B). Analyzing the intersection of disease-associated cell marker genes with otherwise distinct markers from control epithelial cell types, we observe AT*i* cells to share some features with ATI (NTM, GPC5), ATII (SERPINA1, SFTA2) and Aberrant Basaloid cells (Fig. 1C,D). The distinct combination of positive and negative features characterizing AT*i* cells (Fig. 1D) indicate that these are a unique population of cells straddling the phenotypic boundary of otherwise distinct cell states in health and disease. AT*i* cells share only a subset of features with Aberrant Basaloid cells (Fig. 1D): Overlapping features include DNA damage-related genes MDM2 and GDF15; elevated levels of TGFβ-activating integrin subunit ITGB6 and distinct expression of TGFβ signaling genes TGFBR2, SMAD4 and MIR100HG - a host gene for micro-RNA let-7a; hedgehog signaling repressor PTCHD4; the Angiotensin-1 converting protease CPA6, and CTSE – a metastatic hallmark of pancreatic ductal adenocarcinoma and Barretts’ Esophagus^19,20^. In contrast, AT*i* cells lack basaloid features TP63 and KRT17; senescence hallmarks CDKN2A and CDKN2B; TGFβ-inducible NOX4, EPHB2, CASC15 and TGFBI; extracellular matrix (ECM) remodeling genes COL1A1, COL5A1, PRSS2, SPINK1 ADAMTS9, MMP7; the epithelial-mesenchymal transition (EMT) hallmark CDH2 and mutation-inducing transposase PGBD5 (Figure 1D).

AT*i* and Aberrant Basaloid cells with the same features are also observed in publicly available datasets of end-stage COVID-19 lungs^10,21,22^ (Supp. Fig. S2A, Supp. Table S8) extending their relevance from chronic to acute etiologies of pulmonary fibrosis.

### Composition of Abnormal Epithelial Cells in IPF

The median percent of epithelial cells classified as Aberrant Basaloid are similar among IPF samples across all 3 datasets, ranging from 2.72% to 2.91% (Figure 1E). AT*i* cells are significantly enriched in IPF samples vs controls in snRNAseq (Wilcox pVal=2.165E-5) and in Adams *et al.* (Wilcox pVal=0.043), though not in Habermann *et al.* (Wilcox pVal=0.22; Fig. 1E). The proportion of AT*i* cells is significantly higher in IPF snRNAseq samples (median 3.87%) when compared to IPF samples from either scRNAseq datasets (1.32%, p=0.0015 in Adams *et al.*; 2.17%, p=0.038 in Habermann *et al.*; Figure 1E), suggesting that cell dissociation survivor bias negatively impacted the recovery of these cells in previous studies^23^.

We explored the relationship between tissue surface density, a proxy for fibrosis, and the percent of abnormal epithelial cells. The decline in surface density across IPF tissues is significantly associated with a decline in the proportion of ATII cells (p=5.34e-4) alongside an increase in large airway-associated secretory (p=0.013) and ciliated cells (p=0.023) (Figure 1F; Supp. Table S2). Neither Aberrant Basaloid nor AT*i* cell proportions had significant relationships with tissue surface density in IPF (Fig. 1F, Supp. Table S2). This suggests that the emergence of these cells pre-dates the development of pathophysiologic characteristics of fibrosis and it is thus a potential target of anti-fibrotic drugs.

Assessing co-occurrence patterns across all lung cores, the most positive compositional correlates of surface density are unsurprisingly ATI and ATII cells (Figure 1G). Notably, the strongest abundance correlates of AT*i* cells are Aberrant Basaloid cells (Spearman’s rho=0.813) followed by epithelial cells in cell cycle (Spearman’s rho=0.442; Figure 1G), indicating spatial associations with both senescent and proliferating epithelial cells.

### The Continuum of States Between ATI, ATII and Aberrant Basaloid Cells

After evaluating disease-relevant cell types in discrete terms, we assessed their phenotypic connectivity to healthy cell types to model continuous trajectories. Common epithelial cell types were re-clustered into granular sub-populations; inter-connectivity between sub-populations was evaluated with the lineage reconstruction technique “partition-based graph abstraction” (PAGA)^24^. Using an edge confidence threshold > 0.2, all healthy cell types remain connected with a plausible lineage topology, and Aberrant Basaloid cells are exclusively connected to the graph via IPF AT*i* cells towards either ATI or ATII cells (Fig. 2A). To avoid spurious connections in trajectory analysis, low confidence edge pruning from the PAGA graph was extended to connections in the original nuclei-level graph, and a force-directed graph layout of pruned nuclei similarities was built to complement pseudotime analysis (Fig. 2A). Lacking certainty in the orientation of trajectories, we chose to model independent trajectories from either ATII or ATI nuclei towards disease-restricted Aberrant Basaloid nuclei. Generalized additive models (GAMs) of genes were fit to changes in pseudotime across nuclei from each trajectory and organized pseudotemporally by expression peak.

**Figure 2:**
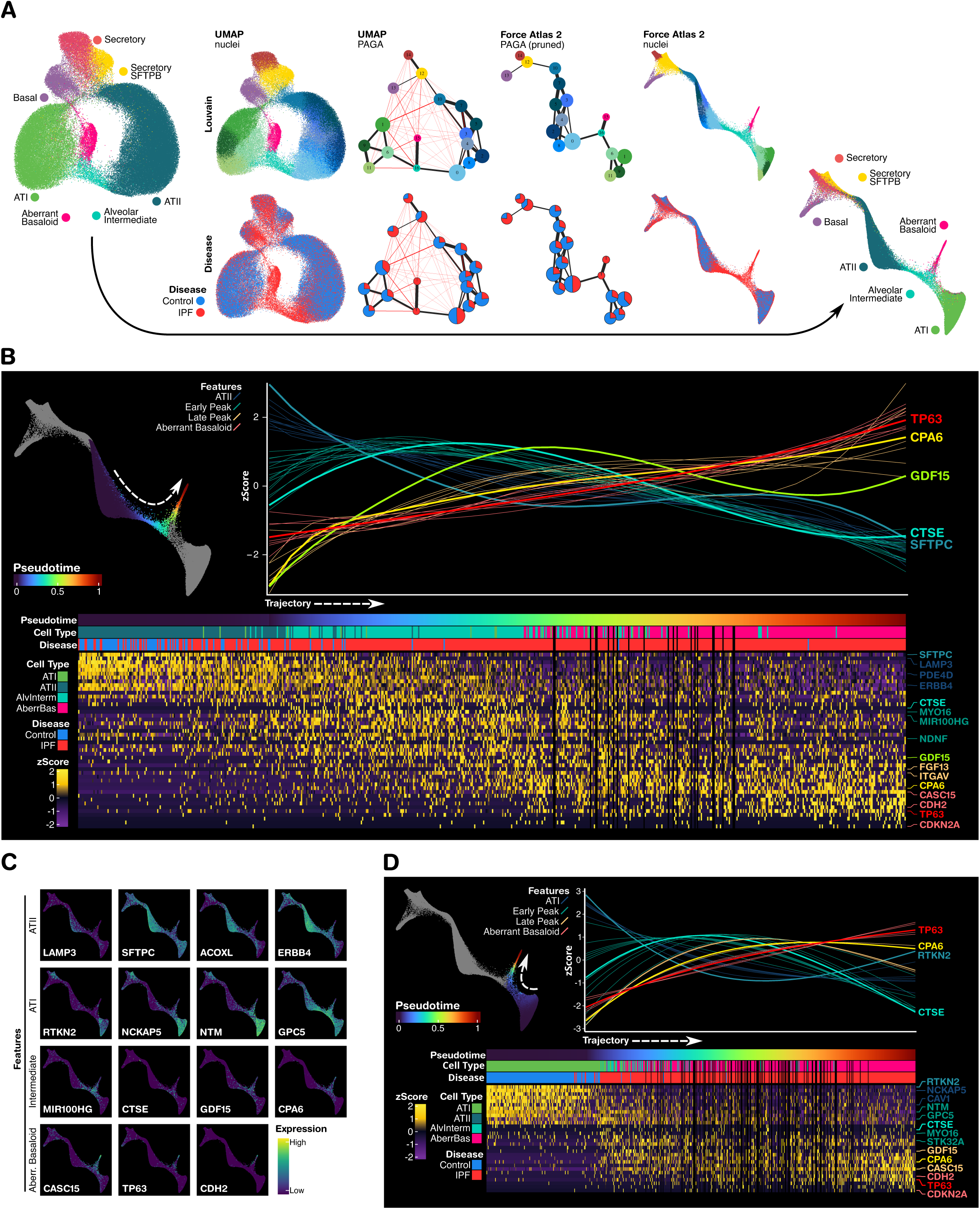
Continuous changes between healthy and disease alveolar epithelial cells. A) From left to right: UMAPs of epithelial nuclei labelled by cell type, Louvain cluster and disease; PAGA graph of Louvain clusters with edges colored by confidence levels above (black) or below (red) the threshold of 0.2; pruned PAGA graph in Force Atlas 2 (FA2) layout; FA2 layout of nuclei with corresponding edge pruning labelled by Louvain cluster, disease and cell type. B) Upper left: FA2 layout of pseudotime trajectory from ATII to Aberrant Basaloid. Upper right: Generalized additive model fits of genes changing as a function of pseudotime. Lower: heatmap of corresponding genes; each column is one nucleus randomly sampled from fixed intervals in pseudotime, gaps correspond to pseudotime intervals with missing representation. C) FA2 layouts colored by gene expression. D) Pseudotime trajectory from ATI to Aberrant Basaloid, represented in the same manner as Fig 2B.

Of the two trajectories, nuclei along the trajectory from ATII towards Aberrant Basaloid features a more gradual pseudotime mapping and more detailed model of pseudotemporal gene dynamics^25^ (Fig. 2B). While markers of ATII cell identity (SFTPC, LAMP3, PDE4D, ERBB4) decline, “Early Peak” features emerge including Let-7a host gene MIR100HG, unconventional myosin MYO16 and the peptidase CTSE (Fig. 2B,C). “Early Peak” features then decline with the loss of cell identity and rise of “Late Peak” genes – including: GDF15, FGF13, ITGAV and CPA6 – before specific Aberrant Basaloid hallmarks emerge (TP63, CDH2, CASC15, CDKN2A; Fig. 2B,C). Trajectory analysis of ATI towards Aberrant Basaloid revealed a similar – though less detailed – sequence of gene expression patterns. ATI features (RTKN2, NCKAP5, CAV1) decline while “Early Peak” features (CTSE, MYO16, STK32A, NTM, GPC5) rise prior to “Late Peak” features (GDF15, CPA6), followed by peaks of Aberrant Basaloid hallmarks (CDH2, TP63, CDKN2A; Fig. 2C,D). Though it remains unclear whether these trajectories represent real lineage fates, this analysis demonstrates that AT*i* cells are a diverse population occupying a branching continuum of phenotypes between ATI, ATII and IPF Aberrant Basaloid cells.

Immunofluorescent microscopy of control and IPF tissues confirmed the presence of both SFTPC+ ATII cells and AGER+ ATI cells with “Early Peak” AT*i* marker CTSE in affected regions of IPF lungs (Fig. 3B) but not in control lungs (Fig. 3A). Despite “Late Peak” features like CPA6 transcriptionally emerging with the loss of cell-type markers (Fig. 2B,D), we observed many SFTPC+/CPA6+ and AGER+/CPA6+ cells in IPF lungs, as well as in hyper-keratinized SFTPC-/AGER-/CPA6+ cells (Fig. 3D), confirming the presence of AT*i* populations in IPF lungs.

**Figure 3:**
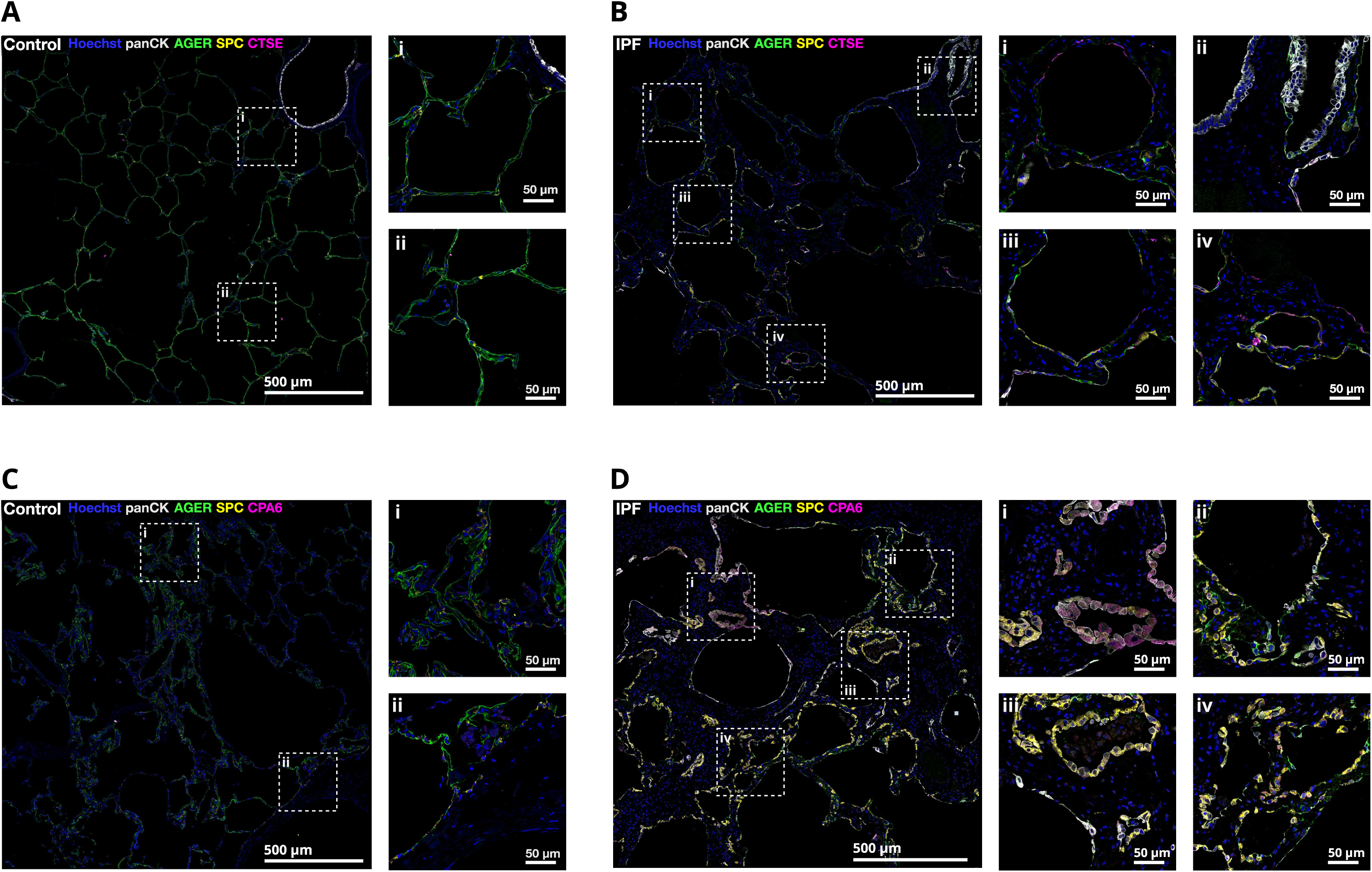
Immunofluorescent Imaging of AT*i* Cells in IPF. A) CTSE expression in control alveolar epithelial cells. B) CTSE expression in IPF alveolar epithelial cells. C) CPA6 expression in control alveolar epithelial cells. D) CPA6 expression in IPF alveolar epithelial cells.

### Longitudinal Assessment of *ex vivo* Fibrosis

To study the emergence of Aberrant Basaloid cells in real time and compare *ex vivo* outcomes to *in vivo* disease, we took advantage of a recent observation that Aberrant Basaloid cells arise *ex vivo* from disease-free human precision cut lung slices (PCLS) following 5 days of exposure to a “fibrotic cocktail” (FC: PDGF-αβ, TGF-β, LPA, and TNFα)^26–28^. We longitudinally assessed this model with snRNAseq (Fig. 4A,B) and observed that both cocktail and control PCLS undergo a decline in epithelial cells over the first 48 hours, before gradually rising (Fig. 4C, Supp. Table S3, Supp. Table S4). Concurrent with this gradual rise is a dramatic increase in epithelial cell proliferation: 3-8% of epithelial cells from 72 hours onwards are undergoing mitosis (Fig. 4C). Hereby, FC exposure significantly reduces the rate of epithelial mitosis between 72 and 120 hours (t-test pVal= 0.010607), from an average of 6.3% in controls to 3.8% with FC (Fig. 4C).

**Figure 4:**
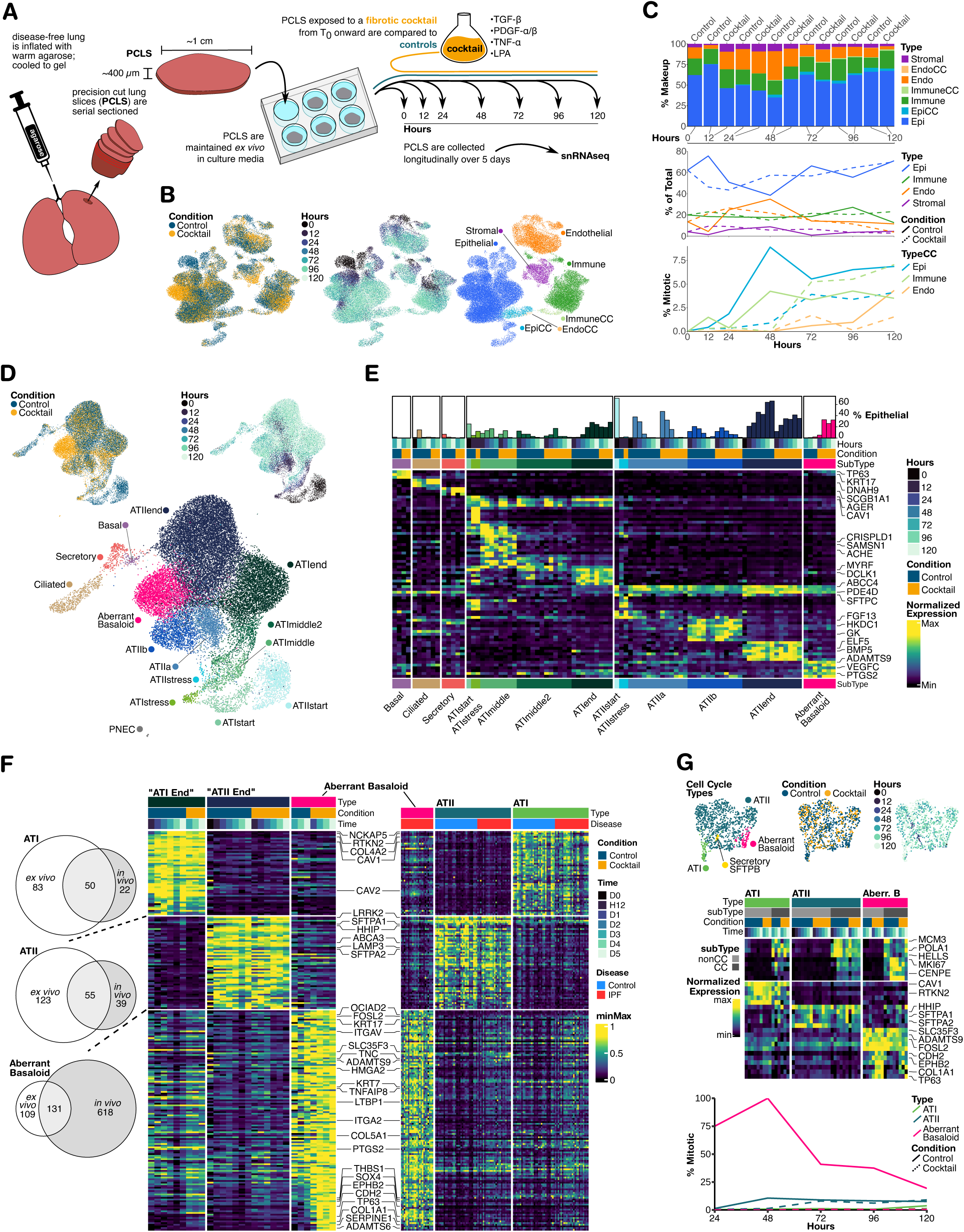
Longitudinal snRNAseq Assessment of PCLS. A) Overview of experiment design. B) UMAP of snRNAseq data labeled by condition, timepoint and cell type. C) Upper: Stacked bar plots of cell type composition. Middle: Line plot of cell type makeup changes as a function of time. Lower: Line plot of the percent of mitotic cells per cell type as a function of time. D) UMAP of epithelial cells labeled by condition, timepoint and cell subtype. E) Heatmap of cell subtype markers and corresponding percent makeup values. The average normalized gene expression value for each sample and subtype are normalized between 0 and 1 across samples. The corresponding percent makeup value is shown in a bar graph above. F) Heatmap of intersecting hallmarks between *ex vivo* and *in vivo* cells. G) Above: UMAPs of epithelial cells in cell cycle labeled by cell type, condition or timepoint. Middle) Heatmap of mitosis marker genes (above) and cell type marker genes for cells undergoing mitosis in cell cycle (CC) or not undergoing mitosis (nonCC). Below: Line plot of the proportion of cells found in cell cycle for each epithelial cell type.

The phenotypic landscape of PCLS alveolar epithelial cells is characterized by many distinct phenotypic substates, associated with different points in time (Fig. 4D,E). All substates could be classified under *in vivo* conventions based on the conservation of marker genes: CAV1, COL4A1, COL4A2 in ATI and PDE4D, ACOXL and ABCC4 in ATII cells (Fig. 4E, Supp. Fig. S3A). The majority of different substates are associated with samples from the first 48 hours, reflecting a period of dynamic behavior with characteristic expression patterns. Early timepoint ATII subtypes exhibit features of stress (HSPA1A, STIP1) and metabolic change (ALDH1L2, GK, KYNU) (Fig. 4E, Supp. Fig. S3E). ATI substates enriched in the first 48 hours exhibit hallmarks of stress (FOS, JUN, HSPA1A), cell cycle arrest (CDKN1C, CCN2, ID2), endoplasmic reticular stress (ERO1B, CLPTM1L, PLA2G4) and cytoskeleton regulation (FSCN1, SAMSN1) (Fig. 4E, Supp. Fig. S3B). PCLS epithelial cells from 72 hours onwards are primarily classified into just three phenotypes: “ATIend”, “ATIIend” and Aberrant Basaloid – suggesting that some degree of homeostasis has been achieved. Of interest, Aberrant Basaloid cells were detected in both control and FC PCLS as early as hour 24 onwards. Aberrant Basaloid cells are significantly more common in FC samples from 72 hours onwards vs control (tTest pVal=0.004385), increasing from an average 1.25% of epithelial cells in control PCLS to 30.4% following FC treatment (Fig. 4E).

A detailed description of temporal ATI and ATII gene expression dynamics is provided in the supplement (Supp. Discussion; Figs. S3, S4). These results depict a biphasic response of alveolar epithelial cells to PCLS experiment conditions where 48 hours of injury response and acclimation are followed by several days of phenotypic stability and tenacious regeneration.

### Comparing control, fibrotic cocktail and IPF Aberrant Basaloid cells

To model the etiology of Aberrant Basaloid cells in PCLS, graph connectivity and pseudotime analysis was performed as described earlier. Similar to *in vivo* results, the strongest connectivities to PCLS Aberrant Basaloid cells are substates of both ATI and ATII cells (Supp. Fig. S3B). Unlike the *in vivo* results, the AT*i* gene signature was not particularly representative of any cell population detected in PCLS, with the prominent exception of FRMD5 and MIR100HG in the PCLS pseudotime trajectory from ATII towards Aberrant Basaloid cells (Supp. Fig. S3E).

Comparing markers of PCLS “ATIend”, “ATIIend” and Aberrant Basaloid to their *in vivo* IPF equivalents, we find most *in vivo* ATI and ATII hallmarks are conserved *ex vivo* (Fig. 4F, Supp. Table S5). Although only 17.5% of the *in vivo* Aberrant Basaloid signal overlaps with ex vivo variants, this overlap encapsulates 54.6% of their defining ex vivo features (Fig. 4F). Control PCLS Aberrant Basaloid cells share many features with FC and *in vivo* variants, including KRT17, ADAMTS9, SLC35F3, PTGS2, HMGA2, FOSL2 and elevated levels of KRT7 and ITGAV (Fig. 4F). However, unlike FC PCLS Aberrant Basaloid cells, control PCLS variants notably lack core disease hallmarks of aberrancy (CDH2, COL1A1, SOX4, SERPINE1, EPHB2) and basal-likeness (TP63) (Fig. 4F). Though Aberrant Basaloid-like cells are observed in control PCLS, FC stimulation induces a more complete molecular representation of Aberrant basaloid cells described in IPF.

Among proliferating epithelial cells, ATII were the most commonly detected, though SFTPB+ Secretory, Aberrant Basaloid and even ATI cells with mitotic hallmarks are found. Strikingly, 30% of all control Aberrant Basaloid cells are observed undergoing mitosis compared to just 0.5% of FC stimulated variants (Fig. 4G); for reference, 7% of all ATII cells and 1.2% of ATI cells are observed mitotic from 24 hours onwards. From hour 72 onwards, FC induced a dramatic decline in the percent of mitotic Aberrant Basaloid (tTest pVal=0.04077) from a per-sample average of 32.6% in controls to just 0.49% (Fig. 4G). FC did not significantly affect the percent of ATII cells undergoing mitosis over this time (tTest pVal=0.6304), indicating that the overall decline in epithelial proliferation induced by FC arose not from direct effects on ATII proliferation, but rather their displacement with senescent Aberrant Basaloid cells.

FC stimulation thus confers a more pathologically relevant presentation of Aberrant Basaloid cells in PCLS in terms of abundance, molecular signature and cell cycle arrest. The observations that control PCLS aberrant basaloid cells are rare yet frequently found dividing, suggest they are normally a transient phenotype associated with homoeostatic repair response. This contrasts with abundant yet rarely dividing Aberrant basaloid Cells found in FC-stimulated PCLS, patterns indicative of a change towards persistence and senescence. Beyond highlighting FC’s ability to emulate the disease phenotype of Aberrant Basaloid cells, these results place a spotlight on control PCLS Aberrant Basaloid cells, understanding their role and plasticity may be key to distinguishing normal repair from maladaptive outcomes.

### Chromatin Accessibility Patterns of Aberrant Basaloid Cells and ATi cells

To explore the epigenetic landscape of epithelial cells in lung fibrosis, we subjected three control and seven IPF lung samples to snMultiOme analysis, where each nucleus is simultaneously profiled with snRNAseq and snATAC-seq chemistry (Fig. 5A). Cell types were classified by transcriptional identity, and ATAC fragments from each cell type per sample were pooled for pseudo bulk peak calling using the R package ArchR^29^; fragments from merged peaks were used in downstream analyses (Fig. 5A). UMAP embeddings of gene expression and chromatin-accessibility show remarkable topological similarities among epithelial cells (Fig. 5B,C). ATI, ATII and Aberrant Basaloid cells occupy unique regions of both RNA and ATAC feature-space. AT*i* straddle the feature-space between these cell types in both UMAP embeddings, however their ATAC profile was far less distinct (Supp. Table S6). The tight co-localization of Aberrant Basaloid cells in unsupervised ATAC-space indicates a divergence from normal epithelial cell chromatin accessibility.

**Figure 5:**
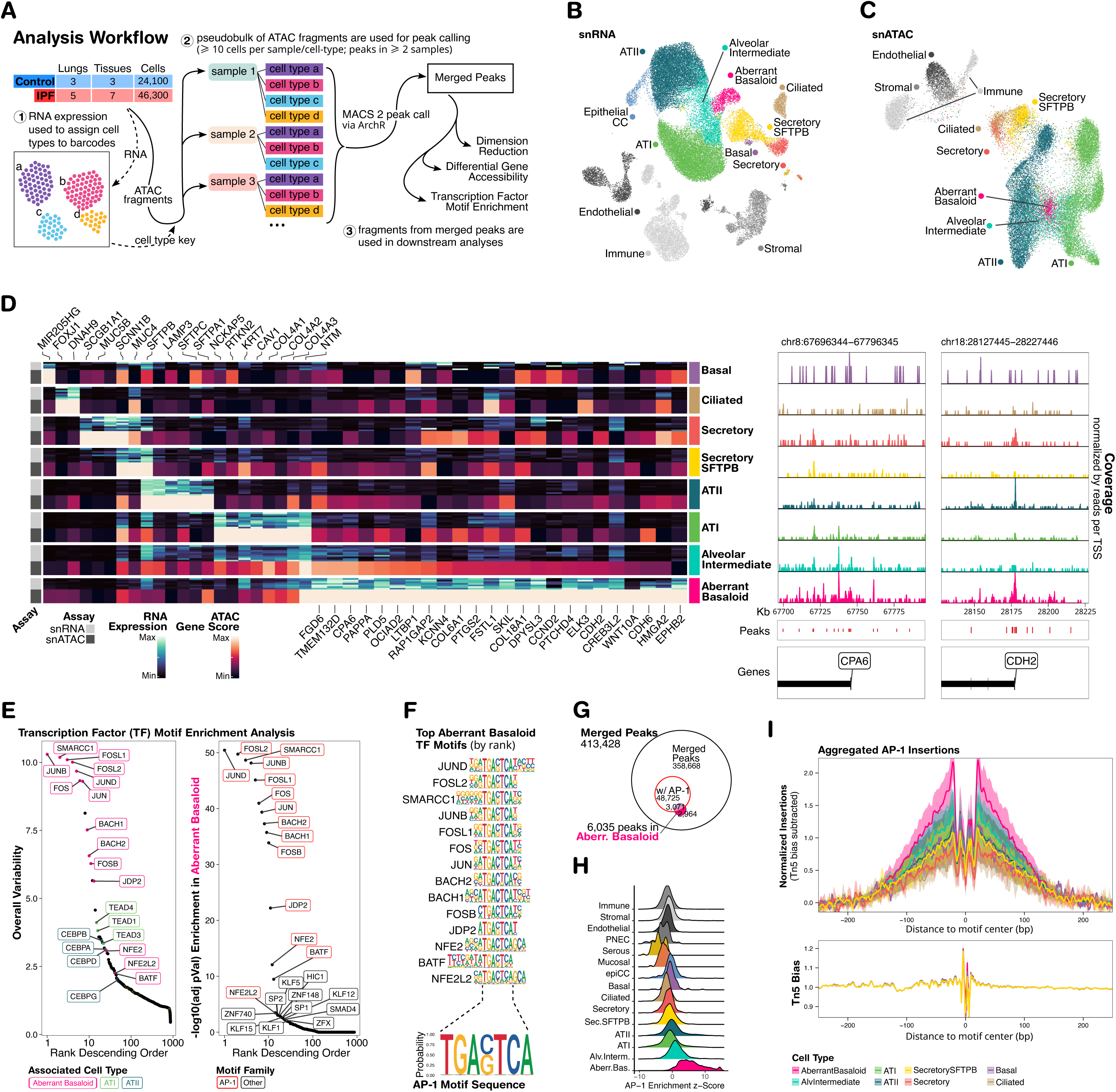
snMultiOme Analysis of IPF and Control Lung Tissue. A) Overview of analysis strategy. B) UMAP of snRNAseq data labeled by cell type. C) UMAP of corresponding snATAC-seq data colored by cell type. D) Heatmap of epithelial gene expression markers (blue) with elevated chromatin accessibility geneScores (red). Average gene expression per sample and average cell type geneScore values for each gene are independently normalized between 0 and 1. E) Left: The top transcription factor (TF) motifs ranked by global contributions to variance; TFs associated with a particular cell type are highlighted. Right: The top TF motifs enriched in Aberrant Basaloid cells compared to other cell types. F) Above: Sequence logos of the top TF motifs in Aberrant Basaloid cells. Below: Sequence logo for AP-1 motif. G) Euler plot showing intersections of all merged peaks, merged peaks significantly enriched in Aberrant Basaloid cells and merged peaks containing at least one AP-1 motif sequence. H) Density plots of AP-1 enrichment z-scores per cell, organized by cell type. I) Footprint analysis of AP-1 transcription factor occupation across epithelial cell types.

We compared differentially expressed genes to differentially accessible genes using ArchR’s Gene Score^29^ (Supp. Table S6). Across epithelial cell types, distinct marker genes often have correspondingly distinct accessibility patterns – MIR205HG in basal cells, FOXJ1 in ciliated, LAMP3 and SFTPC in ATII, RTKN2 and NCKAP5 in ATI (Fig. 5D). Likewise, many uniquely expressed genes in Aberrant Basaloid cells are associated with uniquely accessible chromatin, including TGF-B signaling mediators SKIL, FSTL1, LTBP1 and HMGA2, as well as EMT hallmarks CDH2, COL6A1 and COL18A1 (Fig. 5D). AT*i* cells have no significantly distinct ATAC Gene Scores, but – alongside Aberrant Basaloid cells – they share ATI features like KRT7, NTM and collagen IV and show elevated accessibility of pathological features like CPA6 and the cytoskeleton regulators FGD6 and TMEM132D (Fig. 5D). Collectively, Aberrant Basaloid cells are both epigenetically and transcriptionally distinct from normal lung epithelial cells. The chromatin accessibility profile of AT*i* cells is less categorically distinct, consistent with their diverse intermediate transcriptional states.

### AP-1 Transcription Factor Accessibility is a Defining Feature of Aberrant Basaloid Cells

We explored patterns of transcription factor (TF) motif enrichment across differentially accessible peaks using the TF database CIS-BP^30^. TF motifs with the largest contributions to variance in the data are primarily associated with specific epithelial cell types (Fig. 5E). The top TF families associated with ATII cells are C/EBP, NKX-2 and FOXA1/2; in ATI they are TEAD, KLF, GATA (Supp. Table S7) – observations consistent with previous studies^31–33^. Differentially accessible motif positions from these families have cis-regulatory links to over 60% of their respective transcriptional features (Supp. Fig. S4A,B).

TF motifs for JUND, FOSL2, SMARCC1, JUNB, FOSL1, FOS, JUN, BACH2, BACH1, FOSB and JDP2 are outliers in terms of their contribution to overall variance in the data (Fig. 5E, left) and in terms of the significance of their enrichment among Aberrant Basaloid relative to other cells (pVal<1E-20; Figure 5SE, right). Further examination of the top 15 significantly enriched TF motifs among Aberrant Basaloid cells revealed that they all shared a common sequence: TGA G/C TCA (Fig. 5F, Supp. Table S7) – the consensus sequence for AP-1 – we subsequently focused on this particular sequence motif.

Though 12.5% of all merged peaks contain one or more AP-1 consensus sequences, they are found in over 50% of all differentially accessible peaks in Aberrant Basaloid cells (hypergeometric test p-value < 1e-50; Fig. 5G). Cell-level AP-1 enrichment z-scores are profoundly elevated in Aberrant Basaloid compared to other cell types (Wilcoxon test pVal=2.2e-16; Fig. 5I), where the median cell’s AP-1 enrichment zScore is 5.29; Alveolar Intermediate cells have the second highest median zScore of 1.09. Footprint analysis of accessible chromatin surrounding AP-1 sequences reveals a prominent dip at the binding site among Aberrant Basaloid cells (Fig. 5I), indicating AP-1 TF binding. AP-1 TF motif sequences are found in the majority of differentially accessible chromatin in Aberrant Basaloid cells, these cells have the highest breadth of accessible AP-1 binding sites of any cell in the lung and these sites are actively occupied.

## Discussion

This study provides an unprecedented view of human alveolar epithelial plasticity and unearths important context behind the identity of Aberrant Basaloid cells and their role in regeneration and disease. Analyzing differentially affected IPF tissues, we observe similar frequencies of aberrant basaloid cells across disease tissues, irrespective of their level of fibrosis. We uncover AT*i* cells: a diverse spectrum of cell phenotypes between Aberrant Basaloid cells and ATII and ATI cells – we verify their presence across multiple cohorts and diseases, and localize them in fibrotic tissues. Tracking the emergence of these cells in human PCLS, we observe that fibrotic cocktail stimulation reduces the overall rate of epithelial cell mitosis while increasing the abundance of Aberrant Basaloid cells and imparting them with a molecular profile more aligned with human pulmonary fibrosis. Finally, we establish that IPF Aberrant Basaloid cells have a common epigenetic profile which is distinct from all other lung cell types, characterized by extreme levels of AP-1 family TF chromatin accessibility and binding site occupation.

Upon their discovery, the exotic molecular profile of Aberrant Basaloid cells precluded the ability to infer where these cells originated, a limitation that undercut their translational impact^6,7,34^. Not only was it unclear which cells should be prophylactically targeted to prevent Aberrant Basaloid emergence, but interventions aimed at targeting the behavior of Aberrant Basaloid cells lacked a phenotypic compass to orient the desired outcome. The discovery of AT*i* cells in IPF and fibrotic COVID-19 lungs represents the first example of a “missing link” between these pathological cells and normal epithelial lung cell phenotypes in human lung disease. In both *in vivo* and *ex vivo* models of fibrosis, we observe phenotypic intermediate states between Aberrant Basaloid cells and both ATI and ATII cells – however the PCLS intermediate phenotypes are largely different from the AT*i* cells found in human disease. In addition, studies of cells exposed to fibrosis-associated signaling molecules have shown that human – but not mouse - ATII^35^ and Basal cells^36^ are each capable of achieving an Aberrant Basaloid feature set. Overall, this suggests that the Aberrant Basaloid cell phenotype is an example of cellular convergence: a singular state that different cell types acquire upon injury.

A key limitation of this study is its inability to provide evidence regarding the orientation of changes between these cell types, beyond the assumption that Aberrant Basaloid cells emerge from normal epithelial cells. It is further possible that AT*i* are not transitioning at all. In multiple murine models of fibrosis, it is well established that Transitional Krt8+ cells temporarily emerge from ATII before becoming ATI cells ^11–14^; human disease Aberrant Basaloid cells could reflect malfunction in this process. Perhaps the better analogue to transient murine Transitional Krt8+ cells are the human control PCLS Aberrant Basaloid cells, serving conceptually similar roles in epithelial alveolar regeneration – a form of cellular homoplasy.

Genome-wide chromatin accessibility profiling reveals that IPF Aberrant Basaloid cells share a common epigenetic profile distinct from any other cell type in the lung, forming a unique valley in Waddington’s epigenetic landscape model^37^. Their defining molecular hallmarks are often in regions of the genome uniquely accessible to them, denoting a direct relationship between their functional profile and epigenetic programming.

A defining property of the IPF Aberrant Basaloid epigenome is the degree of AP-1 TF accessibility on display. Gene expression of AP-1 is widely associated with cellular responses to injury and stress, including in lung epithelial cells following cigarette smoke exposure^38^, viral^39^ or bacterial infection^40^, or in the stress response of adjusting to the PCLS environment (Supp. Fig. S4B). However, FOSL2 is the only AP-1 family gene we observe constitutively expressed in either IPF or *ex vivo* Aberrant Basaloid variants, although Footprint analysis indicates that TFs are occupying these binding sites. It is conceivable that these changes arose from transcriptional AP-1 responses to injuries which occurred long before the fibrotic lungs described herein were profiled. Although the AP-1 motif has been among many TF motifs enriched in murine Transitional Krt8+ cells^11,12^ and aged human AT2 cells^41^, this is the first time that genome-wide AP-1 epigenetics are implicated in epithelial cell dysfunction of human non-cancerous lung disease.

Similar AP-1 epigenetics are central to the first recognized instance of innate memory in non-immune cells: skin epithelial stem cells with a history of injury exposure maintained enhanced AP-1 chromatin accessibility and sustained AP-1 binding^42,43^. Retaining this inflammation memory allows cells to respond to future injuries faster and more robustly^44^. Maladaptive outcomes of AP-1 injury memory have been implicated in several diseases where chronic injury leads to epithelial stem cell metaplasia: from benign cases like nasal polyps in chronic rhinosinusitis to precancerous conditions like Barrett’s Esophagus and pancreatic intra-epithelial neoplasia^19,20^. Our study describes the pulmonary disease rendition of AP-1 epigenetic associated dysfunction: where abnormal Aberrant Basaloid cells retain the memory of injuries responsible for triggering lung fibrosis.

In summary, our study provides detailed evidence that alveolar epithelial cell plasticity and regeneration play a significant role in human lung disease. Our findings establish maladaptive epithelial plasticity and regeneration as key mechanistic events in the shift of alveolar epithelial cells from injured innocent bystanders to potential drivers of disease, as proposed by Pardo and Selman nearly 20 years ago^3^. Beyond their pathogenetic significance, these observations highlight the restoration of normal epithelial differentiation as a key therapeutic aim in pulmonary fibrosis.

## Data availability

Raw sequencing data generated for this study was deposited at Gene Expression Omnibus (GEO) under the accession number GSE286182.

## Acknowledgments

We are grateful to all patients and control subjects who participated in this study. Special thanks to Mei Zhong from the Yale Stem Cell Center genomics core facility who performed sequencing, and Joerg Niklaus from the Yale West Campus Imaging Core for provided microscope training and guidance.

## Authors Contribution

NK supervised the study. NK and JCS conceptualized the study and acquired funding. JM performed the micro-CT. JCS, AB and TSA performed snRNAseq and analysis. TSA, AB and AJ performed snMultiOme and analysis. TSA performed immunofluorescent microscopy experiments. JK and AJ performed the PCLS time course experiments. AB, TA and JCS analyzed the COVID19 scRNAseq data. LDS, BMV, MZO and WAW assembled the IPF cohort and supervised the biobanking. IR, JCG provided healthy lung tissue. LDS, JD, FN, JCG, BVP, FA, RJH, IR, XY, BMV, WAW provided critical interpretation, review, and commentary on data and the manuscript. The manuscript was drafted by TSA, JCS and NK and reviewed and edited by all authors.

