## Supplement for "Alveolar epithelial cell plasticity and injury memory in human pulmonary fibrosis"

#### Supplement: Detailed Material and Methods

<u>Selection of study cohort and study approval</u>

Tissue samples were used from patients with IPF undergoing lung transplantation as well as donor lungs not suitable for transplantation as controls from the biobank of the lung transplantation program at University Hospital Leuven, Belgium. Human lungs had been collected following local hospital ethical committee approval (ML6385) and informed patient consent. According to Belgian legislation, declined donor lungs can be used for research purposes. A secondary approval at Yale Institutional Review Board was obtained (# 2000025427).

Tissue processing

In total, 54 lung core samples from 9 IPF lungs and 10 donor lungs were analyzed. Tissue processing was performed as previously described[17]. Following transplant surgery, explanted lungs were frozen in the vapor phase of liquid nitrogen after pressure inflation to 30 $cmH_2O$ and maintenance at 10 $cmH_2O$, as previously described[17]. Whole, frozen lungs were imaged using a high-resolution CT scanner (Siemens Somatom). Then, frozen lungs were cut into 2cm-thick discs along the transversal plane. A core drill with a diameter of 1.4 cm was utilized to sample tissue samples systematically. For this study, one core was randomly selected from each of upper, mid, and lower lung regions for 3 samples per lung. Following X-ray microtomography (see next paragraph), tissue cares were cut into 2mm thick serial slices for downstream analysis (single nuclei RNAseq, single nuclei ATACseq, microscopy). One 2mm thick slice each was formalin-fixed and paraffin-embedded for H&E staining.

X-ray microtomography

Frozen lung samples (diameter 1.4 cm and height 2 cm) were scanned on a custom designed cooling stage by X-ray microtomography (microCT; Skyscan 1172, Bruker) as described previously[17]. Image reconstruction was performed using the NRecon software (Bruker) and analyzed using the CTAn software (Bruker) to segment air from air tissue. Tissue percentage (tissue%) and surface area/volume (surface density) were measured per core.

Single nuclei RNA sequencing

To process the tissue samples, small tissue pieces were put into a gentle MACS C-Tube. 2 mL of nuclei lysis solution (2 M Sucrose, 250 mM Citric Acid, NxGen RNAse Inhibitor 40U/ul) was added to the C-Tube, allowing the tissue to thaw while keeping the tube on ice to maintain low temperatures. Using the gentle MACS Octo dissociator, the “m_lung_01” program followed by the “m_lung_02” program was run to dissociate the tissue. Next, the mixture was centrifuged at 500g for 5 minutes at 4°C, carefully discarding the supernatant while keeping the pellet undisturbed. The pellet was resuspended in 2 mL of nuclei wash buffer (2 M Sucrose, 250 mM

Citric Acid, 10% (w/v) BSA, 1M DTT) and centrifuge again under the same conditions (500g, 5 minutes, 4°C). The supernatant was discarded carefully. To clean up the single-nuclei suspension, an iodixanol gradient was used, keep the nuclei suspension on ice at all times. A 5 mL round-bottom FACS tube was prepared by adding 1 mL of a 30% iodixanol in Nuclei Wash buffer cushion solution. The pellet was resuspended in 500 µL of nuclei wash buffer, then add 500 µL of 50% iodixanol in Nuclei Wash buffer, and mix thoroughly. This mixture was filtered through a 100 µm cell strainer, using the plunger of a 3 mL syringe if needed to gently press the mixture through the filter, ensuring no clumps or tissue chunks remain. The filter was rinsed with 200 µL of nuclei wash buffer and mixed well.

Carefully, the filtered nuclei suspension was layered onto the 30% iodixanol cushion in the prepared FACS tube. This layered solution was centrifuged at maximum speed (4696g) for 20 minutes at 4°C. After centrifugation, the supernatant was removed and discarded. The nuclei pellet was resuspended in 1 mL of PBS containing 1% BSA. The suspension was transferred to a 1.5 mL low-bind Eppendorf tube, then centrifuged at 600g for 5 minutes at 4°C. The supernatant was discarded, then another 1 mL of PBS with 1% BSA was added, and centrifuge again under the same conditions (600g, 5 minutes, 4°C). The final pellet was resuspended in 500 µL of PBS with 1% BSA, ensuring thorough mixing to prevent nuclei clumping. The suspension as filtered through a 20 µm filter into a new 1.5 mL low-bind Eppendorf tube, then the nuclei counted.

<u>Single-cell barcoding, library preparation, and sequencing</u>

Around 20’000 nuclei (PCLS) were loaded on a Chip G with Chromium Single Cell 3′ v3.1 gel beads and reagents (3′ GEX v3.1, 10x Genomics). Final libraries were analyzed on an Agilent Bioanalyzer High Sensitivity DNA chip for qualitative control purposes. cDNA libraries were sequenced on a HiSeq 4000 Illumina platform aiming for 150 million reads per library and a sequencing configuration of 26 base pair (bp) on read1 and 98 bp on read2).

Processing Sequencing Data

To minimize technical variance in gene expression between datasets, fastqs from scRNAseq and snRNAseq data – when available - were processed similarly. Base calls were converted to reads with the software Cell Ranger's (v4.0.0) implementation *mkfastq*. Multiple fastq files from the same library and read strand were catenated to individual read1 and read2 files before trimming. The program cutadapt (v3.0) was used to trim and filter reads prior to genome alignment. For all 10X Genomics 3-prime gene expression data (the *in vivo* snRNAseq (v3.1), *ex vivo* PCLS snRNAseq (v3.1), the RNA data from the snMultiome (v1) experiment and the reprocessed scRNAseq from Adams *et al.*[6] (v2)), read2 files were subject to two passes of contaminant trimming (i) for the template switch oligo sequence (AAGCAGTGGTATCAACGCAGAGTACATGGG) anchored on the 5′ end and (ii) for poly(A) sequences anchored to the 3′ end. Data from the Habermann *et al.* dataset (GSE135893) generated with the 10X genomics 5-prime assay (v2), where the first pass removed R2 5-prime bound template switch sequence: AAGCAGTGGTATCAACGCAGAGTACTTTTTTTTTTTTTTTTTTTTTTTTTTTTT while the second pass removed R2 3-prime bound poly(T) sequences. The Habermann *et al.* dataset included one sample processed with the 10X Genomics 3-prime (v2) assay which we discarded to avoid technical variance within the dataset. Following trimming, read pairs were subsequently removed if the read2 was trimmed below 30 bp or if the read1 barcode sequence had more than 1 bp with a phred < 20.

Trimmed reads were subject to genome alignment using the program STAR (v2.7.6a) and its STARsolo implementation. All libraries were mapped to the same STAR genome index of GRCh38.p13 using GENCODE's human release 37 annotation scheme. Cell and UMI barcodes positions and manufacturer's cell barcode whitelists were parameterized accordingly for each dataset. Collapsed UMIs with reads that span both exonic and intronic sequences were used for downstream analyses and the percent of reads unambiguously spliced was retained as cell metadata for downstream quality control. Paired snRNAseq and snATACseq snMultiome data was simultaneously processed using Cellranger (v7.1) per the manufacturer's instructions, though only the STARsolo RNAseq expression values were utilized for gene expression analysis.

snRNAseq and scRNAseq data from publicly released Covid19 studies (GSE171668, GSE171524, GSE158127) were not readily available in fastq form so processed counts were used.

*in vivo* snRNAseq Cleaning

Processed snRNAseq data cleaning, cell type labeling and exploratory analysis were performed in R (v4.3.3) using the package Seurat (v4.4.0). An iterative and recursive process of increasingly granular cluster analyses were performed to clean the data of spurious and low-quality nuclei and generate preliminary cell type annotations. To prioritize intrasample signals over intersample variance, each sample was cleaned independently. Barcodes with at least 200 transcripts were normalized with a scale factor of 10,000 transcripts per cell, then natural log transformed with a pseudo count of 1. The top 2,000 variable genes were scaled across nuclei and used for principal component analysis (PCA); the top 50 PCs were used for a UMAP layout and Louvain clustering. Clusters of low transcript abundance outliers which lacked defining features were flagged as low-information nuclei and discarded. Clusters with features representative of two or more otherwise-distinct populations found in the sample were flagged as heterotypic multiplets and subsequently discarded. i.e. If multiplet cluster A reflects a combination of bona fide clusters B and C, then differential expression of A vs B will return C's features, and A vs C will return B's. Clusters with a relatively high abundance of mitochondrial RNA, a high percent of spliced transcripts and/or extremely low abundances of the common nuclear-specific lncRNA MALAT1 were flagged as cytoplasmic debris and discarded. An exception was made for B cells who underwent class-switch recombination, as STAR's genome alignment classified IGHA and IGHG transcripts as spliced rather than somatically altered.

Clusters with quality nuclei profiles were assigned a cell type label. Similar cell types were subset from the data, and the process of dimension reduction, clustering, cleaning and annotating was repeated until new populations of either bona fide or spurious nuclei could no longer be resolved. e.g. for epithelial cleaning, the first iteration would span all cells, the second would only include epithelial cells, followed by separate iterations for either alveolar epithelial and airway epithelial cells, and so forth.

*in vivo* snRNAseq Data Integration

To reduce unwanted signals from intersample variance, graph embeddings of the *in vivo* snRNAseq data were performed with data integration at the sample-level using Seurat's implementation of reciprocal PCA (RPCA). Reference samples for integration were selected based on the diversity of cell types present and the level of detail/granularity observed during the cleaning process. Graph-based analyses in both Figures 1 and 2 use the same 20 samples as reference for integration; specifically: all 15 core samples from the 5 IPF lungs (204I, 325I, 190I, 319I, 127I; 3 cores each) and 5 samples from 5 control lungs (248_111_D, 274_62_Db, 145_22_D, 304_71_Db, 222_68_D).

Figure 1's integration includes data from two additional datasets. Epithelial cells from the reprocessed data from Adams *et al.*[6] (GSE136831) and Habermann *et al.*[5] (GSE135893) were identified by their originally reported cell type and combined with the epithelial snRNAseq data. During integration, each scRNAseq dataset was considered one sample (i.e. 57 sources of information were integrated: 55 snRNAseq samples and two scRNAseq datasets).

After splitting the data by either dataset or snRNAseq sample, the top 2,500 variable genes were selected from the 20 aforementioned reference samples using Seurat's *SelectIntegrationFeatures* implementation. Eleven of these selected features were heavy or light-chain immunoglobulins and were subsequently dropped. The number of transcripts and the percent of transcripts mitochondrial were both regressed out during the within-sample scaling and PCA steps of RPCA. Seurat's *FindIntegrationAnchors* was run using the aforementioned references the 2,489 filtered features, a *max.features* value of 250 and the *reduction* argument set to "rpca". The final *IntegrateData* implementation was run under default parameters.

Figure 2's integration only includes snRNAseq data from select epithelial cell types. *SelectIntegrationFeatures* was run on the same aforementioned reference samples for 2,000 variable genes, 11 immunoglobulin genes and one mitochondrial gene (MT-CYB) were removed from the selected features. Sample-level scaling and PCA and subsequent usage of Seurat's *FindIntegrationAnchors* and *IntegratedData* were run in the same manner as described for Figure 1.

Gene Expression Marker Comparisons

Epithelial cell type marker genes were estimated using normalized empirical gene expression values, and calculations were performed for each dataset independently using the Seurat function *FindAllMarkers* with a minimum log2 fold change of 0.5.

For the cell type marker analysis described in Figures 1C and 1D, Core Disease markers for Aberrant Basaloid or Alveolar Intermediate cells were defined as genes with an adjusted pValue less than 1e-10, a pct.2 value below 0.4 and avg log2 fold change value greater than 0.9 or 0.6 respectively – in at least 2 out of 3 datasets. Normal epithelial marker genes were defined by recalculating markers in a similar, but only including control samples in each dataset, and only making comparisons between the 5 normal cell types of interest: ATI, ATII, Basal, SecretorySFTPB and Secretory. Core genes were defined as genes with a log2 fold change greater than 1 and adjusted pValue less than 1e-50 in all 3 datasets; genes passing these criteria in more than one cell type were dropped.

For the cell type marker comparison between *in vivo* and *ex vivo* epithelial cells shown in Figure 4F, PCLS populations ATIend, ATIIend and Aberrant Basaloid were used to represent the *ex vivo* comparison, and both IPF and control ATI, ATII and Aberrant Basaloid were compared in the *in vivo* representation. Each comparison was performed with Seurat's *FindAllMarkers* where only genes with a log2 fold change greater than 0.25 were tested. Numbers representative of sensitivity (pct.1) and specificity (pct.2) were used to calculate the diagnostics odds ratio (DOR) for each gene in each cell type, and only genes with a DOR greater or equal to 3 were used to represent each cell type in each comparison.

Cell Composition Statistics

When comparing the composition of epithelial cell types between disease from different datasets (Fig. 1E), only samples with at least 200 total epithelial cells were included; an unpaired Wilcoxon test was used. When assessing the relationship between snRNAseq epithelial composition and tissue density (Fig. 1F), the R implementation lm was used to fit linear models for control and IPF tissue samples independently. Composition tests of PCLS cells were done between T72 and T120 samples (Fig 4C,E,G) with a t-test.

*in vivo* snRNAseq Graph Embeddings and UMAP layouts

Graph embeddings and UMAP layouts for Figure 1 and 2's analyses were generated by taking their respective integrated data values and subjecting them to Seurat's *ScaleData* and *RunPCA* implementations under default parameters. UMAP layout parameters were chosen for optimal clarity and data density.

Figure 1's UMAP was generated with Seurat's RunUMAP implementation with the first 18 principal components, 35 nearest neighbors, min.dist of 0.5, spread of 0.6, repulsion.strength of 0.8, for 500 epochs with a learning rate of 0.5 and the random seed 7. For the final clustering for cell type annotations, the same PCs and neighborhood size parameters were used for Seurat's *FindNeighbors* followed by *FindClusters* with a resolution parameter of 1.5 for 50 iterations with the seed 7. Cell type labels from these renamed clusters were used throughout downstream analysis.

Figure 2's graph embeddings utilize a non-contiguous sequence of principal components: 1-8, 10-11 and 13-14 - PCs 9 and 12 were avoided for representing variance from background RNA signals of endothelial, fibroblast and T cells. Figure 2's UMAP uses these PCs with Seurat's *RunUMAP* and the following parameters: n.neighbors=15, spread=0.6, repulsion.strength=0.8 for 1000 epochs with the learning rate 0.5, seed 7.

*in vivo* Pseudotime Modeling

Our pseudotime analysis strategy begins by embedding nuclei in a graph based on feature space, clustering nuclei in the graph into communities more granular than cell type and assessing connections between these communities with partition-based graph abstraction (PAGA[22]**)**. To impose constraints on pseudotime trajectories, we prune spurious inter-community edges in both the PAGA and nuclei graph, and subject the pruned nuclei graph to a Markov-chain based approach for generating pseudotime values and branch probabilities (Palantir[41]). Lastly, we evaluate changes in gene expression as a function of pseudotime using generalized additive models (GAMs; tradeSeq[42]).

The R package SeuratDisk's (v0.0.0.9019) *SaveH5Seurat* and *Convert* implementations were used to export the R Seurat object from Figure 2A's UMAP to an h5ad formatted file, accessible for analysis with python's (v3.9.16) Anndata (v0.8.0) and Scanpy (1.9.3) libraries. Due to Scanpy's inability to support non-contiguous PC feature selections, PC coordinates and loadings for PC9 were replaced with PC13's, and PC12's were replaced with PC14's prior to h5ad export.

The same PC features used for Figure 2A's heatmap (now, PCs 1-12) were used to generate a new shared nearest neighbor graph with Scanpy's *pp.neighbors* implementation, now with 20 nearest neighbors using the "gauss" method for computing connectivities, random state 7. The new graph was subject to Scanpy's *tl.louvain* clustering implementation with a resolution parameter of 1.2, in order to divide cell populations into more granular, uniformly sized subpopulations. These clusters were used for PAGA analysis via Scanpy's *tl.paga* implementation using model "v1.2", where a confidence cutoff of 0.2 was used to distinguish high from low confidence edges in the PAGA graph.

Low confidence edges between Louvain clusters in the PAGA graph were pruned and a force-directed graph layout (force atlas 2, FA2) of the pruned PAGA network was created with scanpy's *pl.paga* layout parameter "fa". In the underlying nuclei adjacency and distance matrix, corresponding inter-cluster edges were pruned as well. A force directed graph layout (FA2) of the pruned nuclei graph was generated scanpy's *tl.draw_graph* implementation with layout parameter “fa”, for 500 iterations and nuclei coordinates initialized by their respective Louvain cluster’s position in the pruned PAGA FA2 layout.

ATI, ATII, Alveolar Intermediate and Aberrant Basaloid nuclei were subset from the rest of the data for pseudotime. Pseudotime preprocessing was performed with the scanpy external implementation *tl.palantir* with the following parameters: knn=20, n_components=5, impute_data=False, use_adjacency_matrix=TRUE, where the adjacency matrix and distance key correspond to the pruned graph following PAGA.

The emergent nature of Aberrant Basaloid cells in disease motivated us to orient the pseudotime trajectory from healthy epithelial cells towards this disease endpoint. To achieve a convergent trajectory from either ATII or ATI towards Aberrant Basaloid, we created a branching trajectory from Aberrant Basaloid cells towards either ATII or ATI, and later inverted the pseudotime values to reverse the orientation (a barcode’s pseudotime coordinate is a value between 0 and 1; thus inverted pseudotime = 1 – original pseudotime). The root cell among Aberrant Basaloid was the cell with the highest expression of HMGA2, the ATI terminal cell had the highest expression of RTKN2, and the ATII terminal cell was chosen by selecting the first cell (index 0) in Louvain cluster 10. The final pseudotime results were generated with the scanpy external implementation *tl.palantir_results* where early_cell was the Aberrant Basaloid root cell, terminal states were the aforementioned ATI and ATII terminal cells, knn=20, use_early_cell_as_start=True and num_waypoints=2500. Palantir results and graph layouts created in scanpy were exported back into R for downstream analyses.

To model changes in gene expression as a function of pseudotime from either ATII or ATI towards Aberrant Basaloid, we needed hard assignments for each nucleus to either ATII or ATI-borne branches, however Palantir only returns soft branch assignments in the form of probabilities. Cells from Louvain clusters originally labeled as either ATII or ATI would be assigned to their respective branches, Alveolar Intermediate cells with pseudotime values >= 0.55 were assigned to the ATII branch if their probability of ATII assignment was above 0.9, other Alveolar Intermediate cells with pseudotime >= 0.55 were assigned to ATI. Aberrant Basaloid and Alveolar Intermediate cells with pseudotime values < 0.55 were assigned to either of the two branches based on the weighted probability of branch assignment from Palantir with the R stats *rmultinom* implementation and IPF and control ATI, ATII and Aberrant Basaloid nuclei were compared random seed 7.

Nuclei assigned to either the ATII towards Aberrant Basaloid branch or the ATI towards Aberrant Basaloid branch were analyzed independently using the R package tradeSeq (v1.16.0). Cells assigned to either ATI-borne or ATII-borne trajectories were downsampled to balance representation across pseudotime. All pseudotime values are between 0 and 1; nuclei were assigned to one of 1,001 bins of 0.001 increments; bins with more than 100 barcodes were randomly downsampled to 100 with R's sample implementation, random seed 7. Generalized additive models (GAMs) of genes were fit to each downsampled nuclei from each trajectory using tradeSeq's fitGam implementation on the raw counts with a negative binomial family and in a mixed model where pseudotime distance was a fixed variable but variance between different cDNA libraries was treated as random. The 'nknot' parameter for the ATII trajectory was 7 while the ATI trajectory only used a 'nknot' value of 3.

Immunofluorescent staining

Formalin fixed, paraffin embedded tissue was cut at 5uM thickness and mounted on slides. Slides were deparaffinized and rehydrated: 5 minutes in xylene (repeated twice), 5 minutes in 100% ethanol, 5 minutes in 75% ethanol, 5 minutes in 50% ethanol and 5 minutes in PBS, Antigen retrieval was performed with a tris-based solution (Vector Labs, H-3301-250) at 95°C for 20 minutes and allowed to cool to room temperature. Slides were then treated with the autofluorescence quencher (Biotium, #23014) for 10 minutes per the manufacturer's guidelines, then washed in PBS for 5 minutes, twice.

Tissue was exposed to a serum-free blocking agent (Dako, X0909) for one hour. Primary antibodies used for indirect labeling were diluted in the same block agent and incubated overnight at 4°C in a humid slide box. Slides were washed in PBS three times for 5 minutes each. Secondary antibodies were diluted 1:500 in a diluent of 2.5% donkey serum; slides were incubated with secondary antibodies for 1 hour at room temperature in a dark slide box; all subsequent steps shield the tissues from light to protect the fluorophores. Slides are then washed twice in PBS for 5 minutes each. Commercially preconjugated antibodies for direct labeling were diluted in 2.5% mouse serum and incubated for 30 minutes at room temperature to quench unbound secondary antibodies, then incubated on slides overnight in a humid slide box at 4°C.

After washing slides twice in fresh PBS for 5 minutes each, another autofluorescence quenching treatment was performed (Vector Labs, SP-8500-15) for 5 minutes per the manufacturer's instructions. Slides were subject to a final was step in PBS, three times for 5 minutes each. Coverslips were mounted with an aqueous mounting media containing Hoechst 33342 (Invitrogen, P36983) and left to cure overnight at room temperature. Slides were imaged with a Leica Stellaris 8 Falcon confocal microscope.

The following antibodies and dilution ratios were used in this study: panCK (clone: AE1/AE3, Invitrogen, #53-9003-82) at 1:500; AGER (clone A11: Santa Cruz Biotechnology, sc-80652) at 1:200; SFTPC (clone H-8: Santa Cruz Biotechnology, sc-518029 AF647) at 1:200; CTSE (polyclonal: Atlas Antibodies, HPA012940) at 1:200; CPA6 (polyclonal: proteintech, 13604-1-AP) at 1:200.

<u>PCLS Experiment</u>

We performed a time course analysis of human PCLS treated with or without Fibrotic Cocktail [24] (5 ng/ml TGF-β, 240-B-002/CF, R&D Systems; 5 PDGF-AB, PHG0134, GIBCO; 10 ng/ml TNF-α, P06804, R&D Systems; 5 μM LPA, 62215, Cayman Chemical) from D1 to D5. Four PCLS slices from a control donor at day 0 and at D1 to D5 stimulated with or without were washed in cold 1X PBS and snap frozen. Nuclei were extracted using the Nuclei Isolation kit (CG000505, 10X Genomics). Briefly and based on the manufacturer's protocol and reagents, the tissue was dissociated on ice, centrifugated and washed. The pellet was resuspended and cellular debris were removed. Following another centrifugation step, nuclei were resuspended and counted.

ex vivo snRNAseq Graph Embeddings and UMAP Layouts

snRNAseq data from the PCLS experiment was cleaned and annotated in manner similar to what was described for the *in vivo* data; however, background RNA was much less of a factor and thus data integration steps were unnecessary.

The UMAP of all PCLS cells (Figure 4B) was generated by scaling the top 2,000 variable genes will regressing out variance from transcript abundance prior to PCA. The first 30 PCs were used with Seurat's *RunUMAP* with 20 neighbors for 1,000 epochs with a minimum distance of 0.8. The UMAP of epithelial cells (Figure 4D) was generated by scaling the top 2,000 variable genes while regressing out transcript abundance and the percent of transcripts mitochondrial before PCA; the top 50 PCs were used with 25 neighbors and a minimum distance of 0.7 for 1,000 epochs with a learning rate of 0.5, random seed 7.

*ex vivo* Pseudotime Modelling

We use a similar pseudotime strategy to the one described for the *in vivo* pseudotime analysis. A new graph of PCLS epithelial nuclei was created with scanpy's *pp.neighbors* implementation, using the same 50 PCs used for the Seurat UMAP (Figure 4D), 15 nearest neighbors, the method parameter set to 'gauss', random state 7. The most abundant cell types (ATIend, ATIIend and Aberrant Basaloid) were subclustered into 23 more granular communities with *tl.louvain* resolution 1.5, random seed 7; less common epithelial subtypes inherit their community assignment from their subtype classification (Figure 4D). This subclustered graph was subject to scanpy's tl.paga implementation using model parameter 'v1.2'.

Paga edges with a connectivity statistic less than 0.25 were flagged for pruning. Additional edges were pruned for representing unlikely etiological relationships. (Supplemental Figure S3B). For example: both 'ATIstress' and 'ATIIstress' express a similar set of heat shock protein signatures, this leads to adjacencies between these two cell types in the first 24 hours which appear – at best – irrelevant to any cocktail-related effect or the emergence of Aberrant Basaloid cells. Other edges removed came from connections between early ATI sub-populations which were bypassing temporal intermediate populations: e.g. a connection from T0 “ATIstart” directly to the predominantly T48 “ATImiddle2”).

After extending the pruning to the nuclei-level adjacency graph and distance key, the data was split into two subsets: one with ATII and Aberrant Basaloid, the another with ATI and Aberrant Basaloid. Each subset was preprocessed with scanpy extension *tl.palantir* implementation with 15 components, 15 nearest neighbors and the impute_data argument set to ‘False’.

For the ATII trajectory, D0__ACGATCACACGCTTAA was used as a root cell and three terminal states were used to represent “ATIIend” (D5CTL__ACCTACCAGAAGAGCA), “ATII_B” (D5CTL__TTCATTGCATGGACAG) and Aberrant Basaloid (D5FC__AGCTTCCTCGTGACTA). For the ATI trajectory, the same Aberrant basaloid terminal state was used alongside the “ATIend” terminal state D5CTL__AATAGAGGTTTACCTT while D0__TAATTCCGTACTGACT was used as the ATI root cell. Each subset of data had pseudotime and branch probability estimated with *tl.palantir_results* with their respective root and terminus states, where knn=25 and max iterations set to 100.

Prior to fitting GAMs to trajectories, nuclei were downsampled from each trajectory in the same manner described for the in vivo analysis; gene expression counts from each trajectory were subject to tradeSeqs’s *fitGam* implementation with independent GAMs fit to each branch in the trajectory; 6 knots were used for the ATII trajectory and 5 knots were used for the ATI trajectory.

<u>snMultiome Filtering, Peak Calling and Graph Layouts</u>

The RNAseq component of the data was cleaned and annotated in a manner similar to our earlier approaches to snRNAseq; ATACseq quality was not a consideration at the start of this analysis. Cell type identities determined by RNAseq signatures were used for all downstream ATACseq analyses.

Analysis of the snATACseq data was conducted in R primarily using the package ArchR (v1.0.2). ATAC fragments with barcodes from each sample’s snRNAseq analysis were subject to ArchR’s createArrowFiles implementation with a minimum transcript start site (TSS) score per cell of 5, a minimum fragments cutoff of 0 and maximum fragment cutoff of 1e9; arrow files were then used for the *ArchRProject* function under default parameters. Peaks were called by first using the *addGroupCoverages* function with cells grouped by their RNAseq annotated cell type, 5 minimum cells, 500 maximum cells, 3 minimum sample replicates and 10 maximum replicates. Next, *addReproduciblePeakSet* was used with samples further grouped by cell types, 5 minimum cells, peaks from the y and mitochondrial chromosomes dropped and the peaksMethod argument set to "Macs2".

For the RNAseq UMAP (Figure 5B), nuclei were split by sample and integrating using Seurat's RPCA technique, with the 5 IPF samples (127_10, 190_28, 204_26_I, 319_42, 327_10_I) and 2 control samples (222_68_D, 302_11) used as reference. 2,500 variable features were identified from each sample using *FindVariableFeatures,* 2,000 variables from the reference samples were further selected with *SelectIntegrationFeatures.* Both transcript abundance and the percent of transcripts mitochondrial were regressed out during each samples scaling process, *FindIntegrationAnchors* was run with the aforementioned anchor features and reference samples and the 'max.features' argument set to 300. *IntegrateData* was then run under default parameters. The integrated results were subject to Seurat's *ScaleData* and *RunPCA* under default parameters; *RunUMAP* was run with the first 30 PCs, 15 neighbors, a minimum distance of 0.8 and 1,000 epochs.

For the ATACseq UMAP (Figure 5C), data integration between samples was not used. Instead of PCA, an iterative approach to latent semantic indexing was used for single value decomposition with ArchR's *addIterativeLSI* followed by *addUMAP* with 25 neighbors for 1,000 epochs, seed 7.

<u>ATAC Gene Score and TF motif enrichment</u>

ArchR's GeneScore method was used to create weighted gene scores of chromatin accessibility based on how many peaks were found near the genes TSS. Gene-centric Differences in chromatin accessibility between different cell types was estimated using ArchR's *getMarkerFeatures* with the useMatrix argument set to "GeneScoreMatrix" and both "TSSEnrichment" and "log10(nFrags)" in the bias argument.

Transcription factor (TF) motif enrichment analysis was performed with ArchR's *addMotifAnnotations* with the "cisbp" motif set. The *getMarkerFeatures* implementation was used as before but with "PeakMatrix" in the useMatrix argument slot, maxCells set to 1,000 and "wilcoxon" in the testMethod. Differentially accessible peaks between cell types with an FDR below 0.05 and log2 fold change greater than or equal to 1 were used with ArchR's *peakAnnoEnrichment* function to identify which TF motifs were most associated with which cell type.

<u>snMultiOme Transcription Factor Cis-Regulatory Analysis</u>

The same approach was used for ATII, ATI and Aberrant basaloid cells independently. Marker genes for each cell type were defined as genes found significantly different in 2 of 3 *in vivo* datasets. The top enriched TF motifs for each cell type were grouped into families, and positions of these TF motifs were identified in the cell type's respective significantly enriched peaks. For each of these positions, the nearest gene was identified with the R package GenomicRange's (v1.54.1) *distanceToNearest* implementation; if the closest gene was defined as marker gene for that cell type, then this relationship is included in the cell type's alluvial and Euler plot (Supplemental Figures S5A-C).

<u>COVID19 Lung Data Analysis</u>

Preprocessed scRNAseq data from Bharat et al. (PMID: 33257409) and two snRNAseq datasets from Melms et al. (PMID: 33915568) and Delorey et al. (PMID: 33915569) were downloaded from NCBI Gene Expression Omnibus (GEO) under the accession numbers GSE158127, GSE171524 and GSE171668, respectively.

Each of the datasets was curated in R (v 4.0.5) using the toolkit in Seurat package (v 4.1.1) and cells annotated as epithelial were isolated for the downstream analysis. Accounting for the intra-dataset patient-level batch effect, epithelial cells were split based on the patients and integration was done using Seurat's reciprocal principal component analysis (RPCA) method by sequentially adding one dataset at a time. Prior to the integration step, mitochondrial and ribosomal genes were excluded from the feature selection and regressed out during the scaling before performing PCA on combined epithelial cells. Final integration parameter regulating number of neighbors for weighting anchors (k.weight) was adjusted to 70. All other RPCA integration parameters were under the default settings.

Cell types were identified after recursive integration, scaling, dimensionality reduction, UMAP embedding, high resolution clustering and differential gene expression analysis of all cells. After each reintegration step, the presence of all cell types separately observed in each of the datasets was strictly followed where the assignment was based on the unique gene expression pattern across multiple subjects in several datasets. Final UMAP embedding was based on principal components 1-6, 8-14, 16 and 18 which clearly isolated all unique cell types observed in the curation of each dataset separately. UMAP implementation was run with 50 neighbors, a minimum distance of 0.5, repulsion strength and spread parameters both equaling 1 with 1000 iterations and learning rate of 0.2 at seed=7.

### Supplemental Results

#### Temporal ATI and ATII gene expression dynamics

A graph topology and pseudotime analysis performed as described earlier across all time points revealed that both ATI and ATII intermediate populations contributed independently to the Aberrant Basaloid Cell Populations (Supp. Fig. S3,S4) across time points with unique sequences of molecular changes as they converge. The ATII pseudotime trajectory has three distinct endpoints enriched in cells from 72h to 120h: "ATIIend", "ATII-B" and FC-associated Aberrant Basaloid cells (Supp. Fig. S3D,E). All ATII cells transiently express stress markers HSPA1A, HSP1D followed by a decline of *in vivo* features FGFR1, FSTL4 and maintained elevated expression of MIR100HG and FRMD5 – hallmarks of *in vivo* AT*i* cells – when committed to Aberrant Basaloid cells.

The ATI trajectory is relatively simple with a single branch event (Fig. S4A). All ATI cells begin with an elaborate stress response involving expression of metallothionines and FOS and JUN family transcription factors followed by short-lived expression of the cell cycle inhibitor CDKN1C and DNA damage marker H2AX. Next, endoplasmic reticular proteins ERO1B and CLPTM1L are expressed alongside the epigenetic regulator HDAC9 during a prolonged period of growth factor BDNF expression, immediately preceding the branching event towards either "ATIend" or Aberrant Basaloid cells (Fig. S4B). CDH2 expression occurs notably early in the ATI-borne trajectory (Fig. S3B); cross-referencing its expression with ATI hallmarks RTKN2, NCKAP5, COL4A1 and COL4A2 expression in UMAPs (Fig. S3A) indicates that EMT occurs prior to the loss of ATI features.

### Cis-regulation of Aberrant Basaloid Features

Only 33.7% of Aberrant Basaloid markers are among the closest genes to a differentially accessible AP-1 position, consistent with AP-1's role as an effector TF. KLF-family motif positions enriched in Aberrant Basaloid are associated with a higher overall share of Aberrant Basaloid genes (35.1%), and over 68% of Aberrant Basaloid genes associated with enriched AP-1 positions are also associated with enriched KLF-family positions. AP-1/KLF associated genes include mediators of TGF-b signaling: TGFB2, SMAD3, LTBP1 and SKIL; cAMP TFs CREB5 and CREB3L2; other assorted regulators: CRABP2, CUX1, PDGFC, KLF6 and PLAU. Enriched KLF-family motif positions were uniquely linked to important marker genes including CDH2, CDKN2A and KRT17 (Figure S4C).

### Supplemental Figure Legends

#### Supplementary Figure S1: Composition Patterns of snRNAseq and scRNAseq Datasets

Above: Table of total lung and cell counts across three datasets. Middle: UMAPs of all cells from each dataset with cells colored by cell type category. Below: Bar plots of the percent makeup of cell type categories across samples for each respective dataset, organized by either surface density or disease.

#### Supplementary Figure S2: Analysis of Epithelial Cells in COVID-19 Induced Pulmonary Fibrosis

A) UMAPs of epithelial cells from COVID-19 datasets labeled by cell type, disease, dataset and subject. B) Heatmap of cell type markers. Each column is the average normalized gene expression from one subject per cell type. Data for each gene is min-max transformed independently for each dataset.

#### Supplementary Figure S3: PCLS Pseudotime Trajectory Analysis of ATII Cells

A) UMAPs of PCLS epithelial cells labeled by cell sub type (left), time point (middle) and conserved markers of ATII (right, upper) and ATI cells (right, lower). Normalized gene expression for each gene is min-max normalized between 0 and 1. B) PAGA of PCLS epithelial cells following pruning of edges below a Confidence of X. Spurious connections between different cell types were additionally pruned (lower left). C) UMAP overlay of ATI and ATII branching trajectories following PAGA constraints. D) UMAP of ATII pseudotime mapping overlayed with the ATII branching trajectory (upper); pruned PAGA used for ATII pseudotime (lower, left); UMAP of cell branch assignments for ATII and Aberrant Basaloid cells. E) Plots of composition and expression changes as a function of pseudotime; all plots share a common x-axis of pseudotime. From top to bottom: a histogram of cell abundances in control and cocktail samples across pseudotime; density plots of cell abundances from timepoints across pseudotime; generalized additive models of genes to cells assigned to different branches across pseudotime. Each gene's expression is min-max normalized between 0 and 1.

**Supplementary Figure S4: PCLS Pseudotime Trajectory Analysis of ATI Cells**

A) UMAP of ATI pseudotime mapping overlayed with the ATI branching trajectory (upper); pruned PAGA used for ATI pseudotime (lower, left); UMAP of cell branch assignments for ATI and Aberrant Basaloid cells. B) Plots of composition and expression changes as a function of pseudotime; all plots share a common x-axis of pseudotime. From top to bottom: a histogram of cell abundances in control and cocktail samples across pseudotime; density plots of cell abundances from timepoints across pseudotime; generalized additive models of genes to cells assigned to different branches across pseudotime. Each gene's expression is min-max normalized between 0 and 1.

**Supplementary Figure S5: Mapping Enriched TF Motif Positions to Cell Type Marker Genes**

A) From left to right: the top TF motifs enriched in ATII cells; sequence logos organized by TF family; alluvial plot linking enriched motif sequence sites from differentially accessible peaks to their closest gene; Euler plot of ATII marker genes organized by cis-regulatory TF. B) Corresponding analysis of ATI cells. C) Corresponding analysis of Aberrant Basaloid.

### Supplemental Table Legends

**Supp. Table S1. Epithelial Marker Table**

Tab-delimited results of Wilcoxon rank-sum test of each epithelial cell type against the other epithelial, per cell type.

**Supp. Table S2. Linear regression Surface Density vs Epithelial Proportions**

Tab-delimited results of linear regression models of changes in epithelial cell proportions as a function of surface density.

**Supp. Table S3. Lineage percentages of non-proliferating cells in the PCLS experiment**

Percentages of PCLS of cells by major lineage annotation (epithelial, endothelial, stromal, immune) excluding cell undergoing cell cycle, per time point and condition.

**Supp. Table S4. Lineage percentages of cells undergoing cell cycle in the PCLS experiment**

Percentages of PCLS cells by major lineage annotation (epithelial, endothelial, stromal, immune) of cells undergoing cell cycle, per time point and condition.

**Supp. Table S5. Marker table of the comparison of *in vivo* IPF vs *ex vivo* PCLS epithelial cell states**

Tab-delimited results of Wilcoxon rank-sum test of each epithelial cell-type against the other epithelial varieties, comparing *ex vivo* PCLS "ATIend", "ATIIend" and Aberrant Basaloid cells and their *in vivo* IPF equivalents, per cell type.

**Supp. Table S6. snATACseq geneScore table**

Tab-delimited snATACseq GeneScore results of the IPF single nuclei MultiOme experiment, per cell type.

#### Supp. Table S7. Transcription factor motif table

Tab-delimited results of transcription factor motif enrichment analysis, per cell type.

#### Supp. Table S8. Epithelial Covid-19 marker table

Tab-delimited results of Wilcoxon rank-sum test of the Covid-19 object of each epithelial cell type against the other epithelial, per cell type and per dataset.

#### Supp. Data. Raw data to figures

Raw data to figures: IPF epithelial frequencies per subject; PCLS epithelial counts of cells undergoing cell cycle; PCLS epithelial counts of cells not undergoing cell cycle; PCLS lineage proportions of cells undergoing cell cycle; PCLS lineage proportions of cells not undergoing cell cycle; PCLS epithelial subtype proportions.
