## Supplementary material for "Alveolar epithelial cell plasticity and injury memory in human pulmonary fibrosis": Supp. Fig. S

| **lungs** (total cells) | **Control** | **IPF** | **COPD** | **Other ILD** |
|---|---|---|---|---|
| snRNA | **10** (292,438) | **9** (222,550) | - | - |
| Adams *et al.* | **28** (95,054) | **32** (144,214) | **18** (67,889) | - |
| Habermann *et al.* | **10** (31,644) | **12** (57,682) | - | **8** (25,070) |

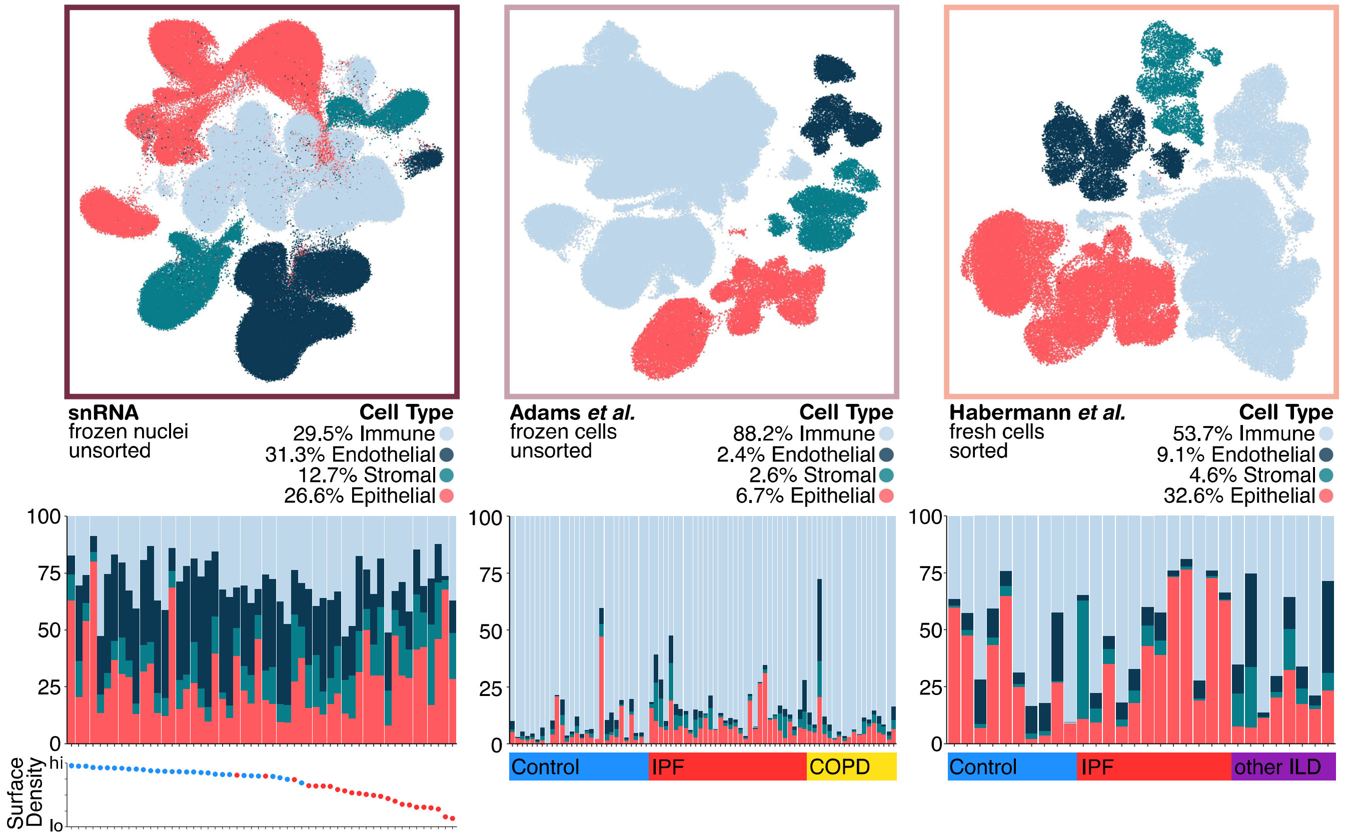


**Supplementary Figure S1: Composition Patterns of snRNAseq and scRNAseq Datasets**
Above: Table of total lung and cell counts across three datasets. Middle: UMAPs of all cells from each dataset with cells colored by cell type category. Below: Barlots of the percent makeup of cell type categories across samples for each respective dataset, organized by either surface density or disease.

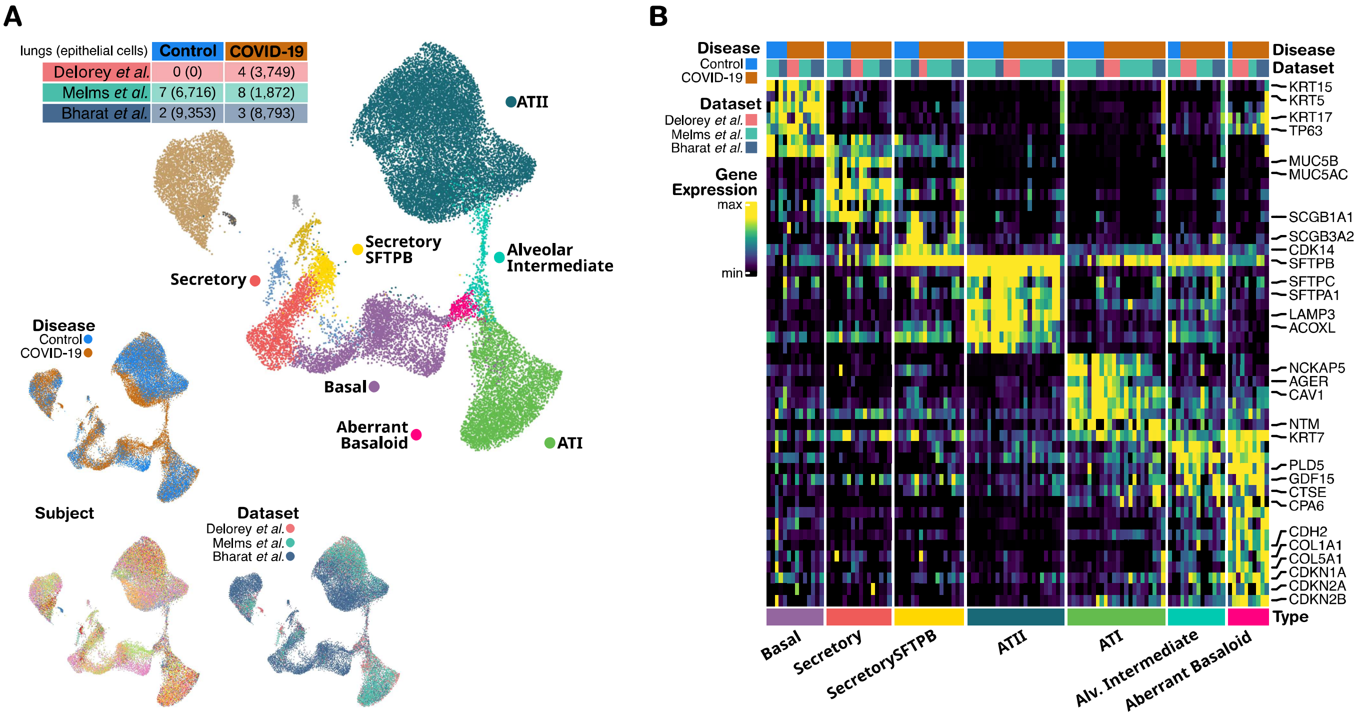


**Supplementary Figure S2: Analysis of Epithelial Cells in COVID-19 Induced Pulmonary Fibrosis**
**A**) UMAPs of epithelial cells from COVID-19 datasets labled by cell type, disease, dataset and subject. **B**) Heatmap of cell type markers. Each column is the average normalized gene expression from one subject per cell type. Data for each gene is min-max transformed independently for each dataset.

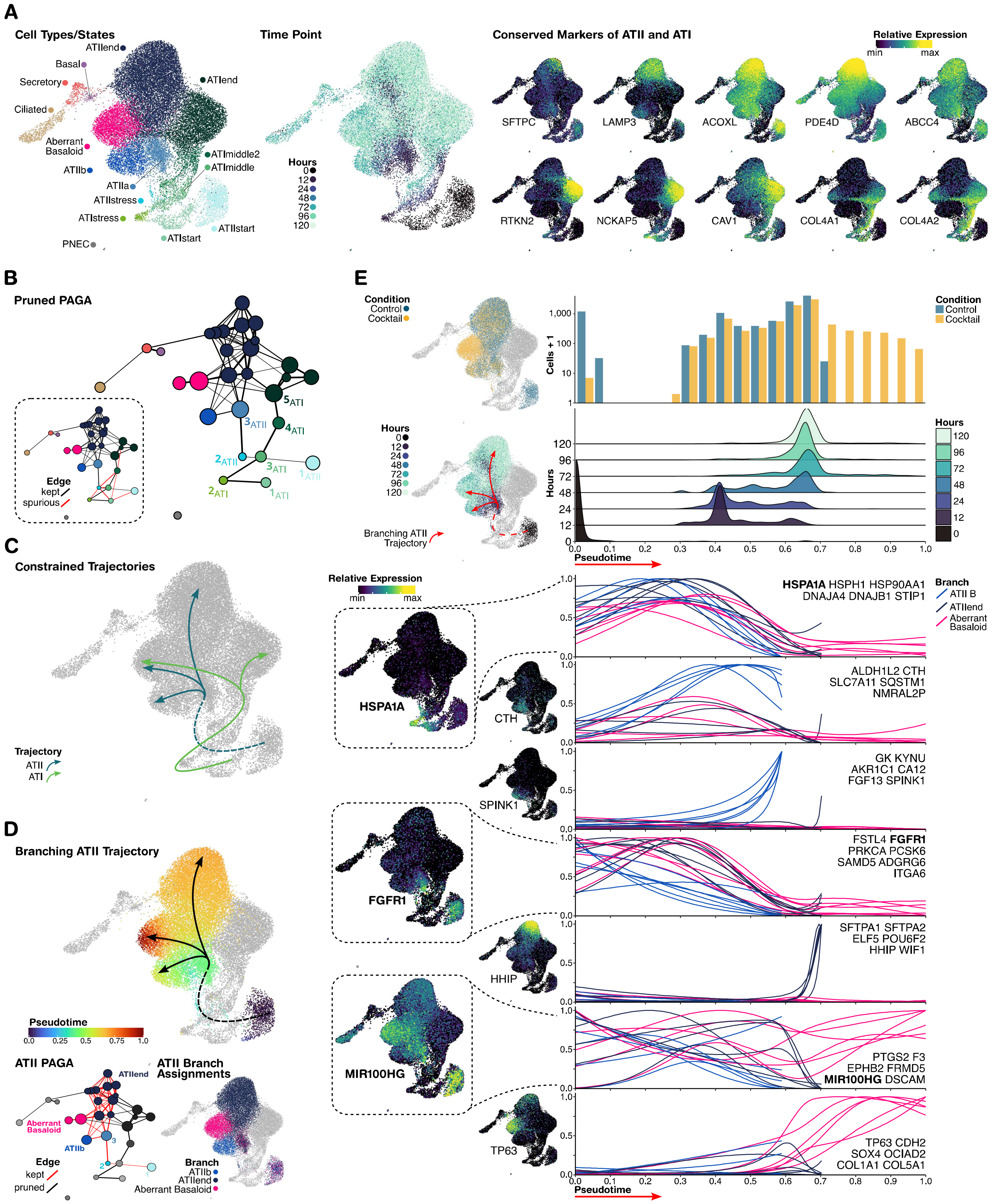


**Supplementary Figure S3: PCLS Pseudotime Trajectory Analysis of ATII Cells**

**A**) UMAPs of PCLS epithelial cells labeled by cell sub type (left), time point (middle) and conserved markers of ATII (right, upper) and ATI cells (right, lower). Normalized gene expression for each gene is min-max normalized between 0 and 1. **B**) PAGA of PCLS epithelial cells following pruning of edges below a Confidence of **X**. Spurious connections between different cell types were additionally pruned (lower left). **C**) UMAP overlay of ATI and ATII branching trajectories following PAGA constraints. **D**) UMAP of ATII pseudotime mapping overlayed with the ATII branching trajectory (upper); pruned PAGA used for ATII pseudotime (lower, left); UMAP of cell branch assignments for ATII and Aberrant Basaloid cells. **E**) Plots of composition and expression changes as a function of pseudotime; all plots share a common x-axis of pseudotime. From top to bottom: a histogram of cell abundances in control and cocktail samples across pseudotime; density plots of cell abundances from timepoints across pseudotime; generalized additive models of genes to cells assigned to different branches across pseudotime. Each gene's expression is min-max normalized between 0 and 1.

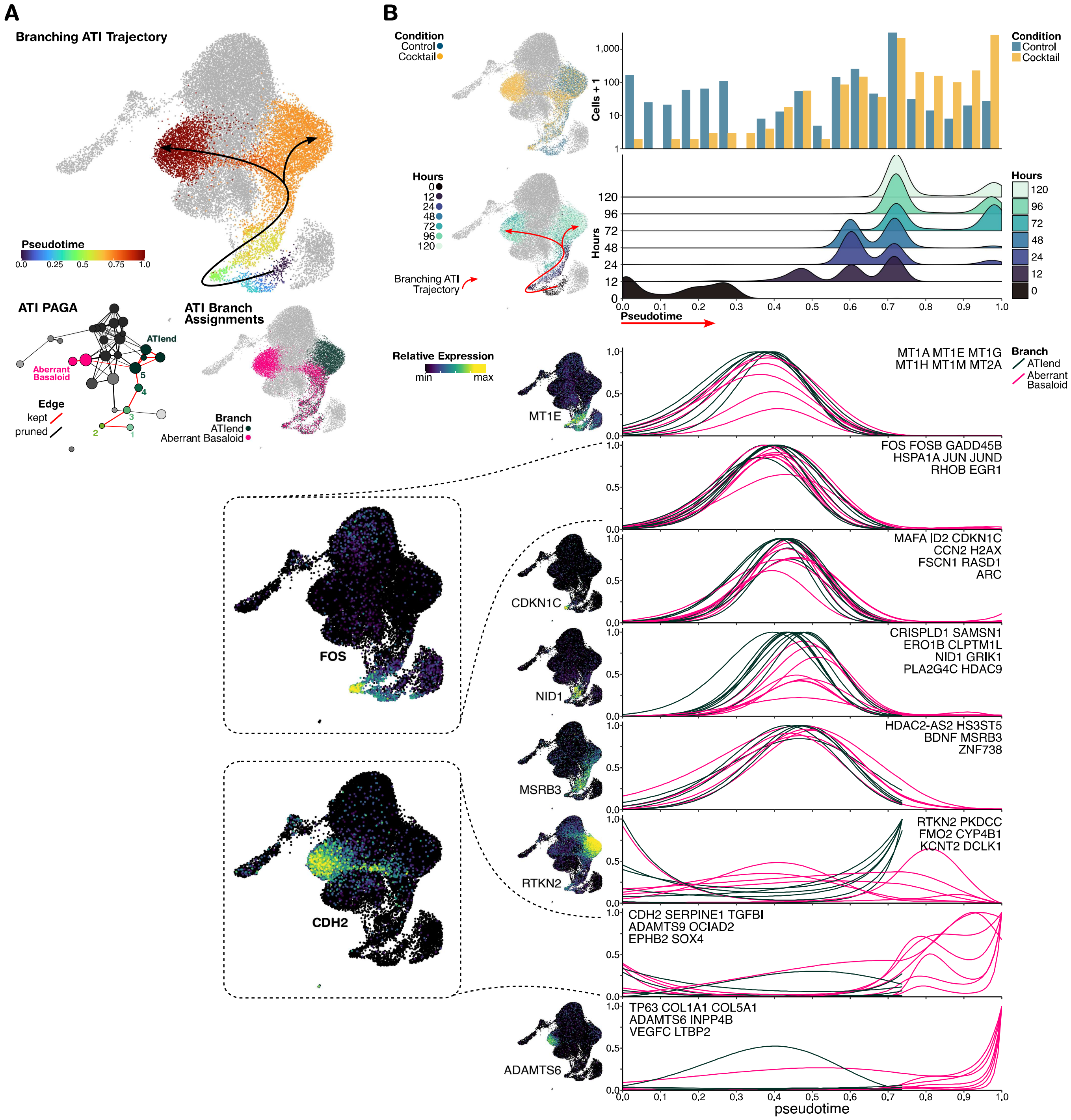


**Supplementary Figure S4: PCLS Pseudotime Trajectory Analysis of ATI Cells**

**A**) UMAP of ATI pseudotime mapping overlayed with the ATI branching trajectory (upper); pruned PAGA used for ATI pseudotime (lower, left); UMAP of cell branch assignments for ATI and Aberrant Basaloid cells. **B**) Plots of composition and expression changes as a function of pseudotime; all plots share a common x-axis of pseudotime. From top to bottom: a histogram of cell abundances in control and cocktail samples across pseudotime; density plots of cell abundances from timepoints across pseudotime; generalized additive models of genes to cells assigned to different branches across pseudotime. Each gene's expression is min-max normalized between 0 and 1.

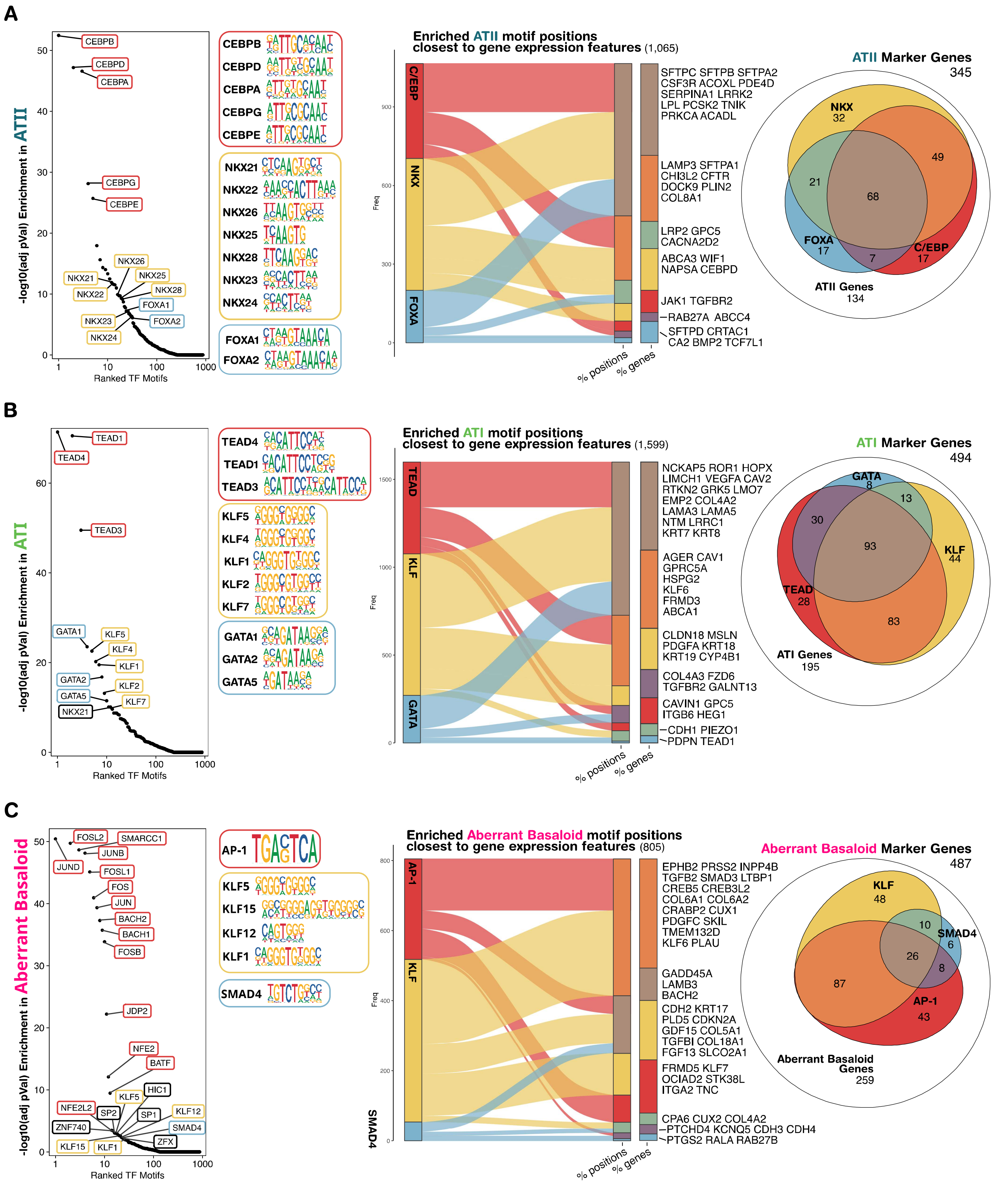


**Supplementary Figure S5: Mapping Enriched TF Motif Positions to Cell Type Marker Genes**

**A**) From left to right: the top TF motifs enriched in ATII cells; sequence logos organized by TF family; alluvial plot linking enriched motif sequence sites from differentially accessible peaks to their closest gene; euler plot of ATII marker genes organized by cis-regulatory TF. **B**) Corresponding analysis of ATI cells. **C**) Corresponding analysis of Aberrant Basaloid.
